# The Statistical Trends of Protein Evolution: A Lesson from AlphaFold Database

**DOI:** 10.1101/2022.04.07.487447

**Authors:** Qian-Yuan Tang, Weitong Ren, Jun Wang, Kunihiko Kaneko

## Abstract

The recent development of artificial intelligence provides us with new and powerful tools for studying the mysterious relationship between organism evolution and protein evolution. In this work, based on the AlphaFold Protein Structure Database (AlphaFold DB), we perform comparative analyses of the proteins of different organisms. The statistics of AlphaFold-predicted structures show that, for organisms with higher complexity, their constituent proteins will have larger radii of gyration, higher coil fractions, and slower vibrations, statistically. By conducting normal mode analysis and scaling analyses, we demonstrate that higher organismal complexity correlates with lower fractal dimensions in both the structure and dynamics of the constituent proteins, suggesting that higher functional specialization is associated with higher organismal complexity. We also uncover the topology and sequence bases of these correlations. As the organismal complexity increases, the residue contact networks of the constituent proteins will be more assortative, and these proteins will have a higher degree of hydrophilic-hydrophobic segregation in the sequences. Furthermore, by comparing the statistical structural proximity across the proteomes with the phylogenetic tree of homologous proteins, we show that, statistical structural proximity across the proteomes may indirectly reflect the phylogenetic proximity, indicating a statistical trend of protein evolution in parallel with organism evolution. This study provides new insights into how the diversity in the functionality of proteins increases and how the dimensionality of the manifold of protein dynamics reduces during evolution, contributing to the understanding of the origin and evolution of lives.

## Introduction

The evolution of organisms takes place over time, as Darwin noted in his *On the Origin of Species*. It is usually naively discussed that the complexity of life has increased throughout evolution, whereas its validity is not always confirmed (McShea 1996; Adami et al. 2000; Furusawa and Kaneko 2000). Generally, it is widely accepted that complexity should be characterized as the difficulty of reducing a system to constitutional components. If the system consists of more different internal components, the complexity is generally higher. According to this idea, one can simply assume that eukaryote cells are more complex than prokaryotes, and multicellular organisms with more distinct cell types are more complex than unicellular organisms, as is also adopted in *The Major Transitions in Evolution* (Maynard Smith and Szathmary 1997). Parallel to the increasing complexity of organisms, proteins, as the basic building blocks of organisms, are also undergoing continuous evolution. Previously, there were evolutionary studies elaborating on the phylogenetic analysis of proteomes (Gerstein et al. 1994; Caetano-Anollés et al. 2009; Caetano-Anollés et al. 2021) or the proteins that share the common ancestral sequence or structure (Labas et al. 2002; Pin et al. 2003; Zardoya 2005; Morcos et al. 2011; Finnigan et al. 2012; Espada et al. 2015). To uncover the connection between organism evolution and protein evolution (Liu and Rost 2001; Koonin et al. 2002; Pál et al. 2006; Zeldovich and Shakhnovich 2008; Sikosek and Chan 2014), combining the view of evolution from two perspectives may be necessary. In the microscopic view (molecule level), the evolution of a protein does not necessarily follow the evolutionary path as the species evolve (Choi and Kim 2006); while in the macroscopic view (organism level), if considering an ensemble of thousands of proteins within the same organism, just as thermodynamic laws and collective order can emerge from the random motions of molecules, it may be possible to discover a “collective” trend consistent with the increase of organismal complexity.

Notably, the recent development of artificial intelligence (AI) provides us with new and powerful tools to help us elucidate the trends in protein evolution on the macroscopic scale. AlphaFold, an AI system developed by DeepMind, which makes full use of the coevolutionary information to predict protein structure, has already won an unprecedented and overwhelming success in protein structure predictions (Senior et al. 2020; Jumper et al. 2021). Despite the limitations noted in previous research (Bagdonas et al. 2021; Pak et al. 2021; Ruff and Pappu 2021), AlphaFold 2 is acknowledged to provide high-accuracy predictions of protein structures, even for sequences with relatively fewer homologous sequences (Jumper et al. 2021). The exceeding accuracy and the speed of AlphaFold 2 provide the possibility of generating an extensive database of structure predictions. AlphaFold Protein Structure Database (AlphaFold DB) provides now provides free access to about 200 million predicted protein structures, which covers the complete proteome of various organisms ranging from bacteria, archaea, unicellular and multicellular eukaryotes to humans (Varadi et al. 2022), and it keeps expanding. AlphaFold DB not only offers the potential to answer critical questions in medical and biological sciences (Robertson et al. 2021; Bayly-Jones and Whisstock 2022), but also shows new possibilities in the study of protein evolution. Instead of focusing only on specific families of proteins (Bayly-Jones and Whisstock 2022), we can now perform comparative structural analyses for the proteins in different organisms. Combining with other essential evolutionary analyses, the statistics of the full-proteome protein structures may help us uncover the hidden connection between protein evolution and organism evolution.

In this work, based on the AlphaFold-predicted structures of the proteomes of 48 organisms, we perform comparative analyses of the constituent proteins of different organisms. The statistical results indicate a correlation between the flexibility of constituent proteins and the complexity of organisms; that is to say, for organisms with higher complexity, their constituent proteins will have larger radii of gyration, higher coil fractions, and slower vibrations, statistically. By conducting normal mode analysis and scaling analyses, we demonstrate that higher organismal complexity correlates with lower fractal dimensions in both the structure and dynamics of the constituent proteins, suggesting that higher functional specialization is associated with higher organismal complexity. We also uncover the topology and sequence bases of these correlations. As the organismal complexity increases, the residue contact networks of the constituent proteins will be more assortative, and these proteins will have a higher degree of hydrophilic-hydrophobic segregation in the sequences. Furthermore, by comparing the statistical structural proximity across the proteomes with the phylogenetic tree of homologous proteins, we show that, statistical structural proximity across the proteomes may indirectly reflect the phylogenetic proximity of homologous proteins among organisms. Such a result suggests a statistical trend of protein evolution in parallel with organism evolution. This study provides new insights into how the diversity of protein functionality increases and how the dimensionality of the protein dynamics manifold reduces in the evolution, contributing to the understanding of the origin and evolution of lives.

## Results

### The statistics of AlphaFold-predicted structures indicate a correlation between the flexibility of constituent proteins and the complexity of organisms

We perform a comparative analysis of the predicted protein structures of the 48 organisms from the AlphaFold DB. The details of the full dataset are listed in Supplementary Information (SI) Table S1. Here, let us first focus on the proteins from the 16 model organisms with similar chain lengths. Namely, for the proteomes of various organisms, we always select the protein structures with chain lengths *N* ≈ 250, calculate their structural characteristics, and conduct statistical analyses. As shown in Fig. 1A, despite having similar chain lengths, the constituent proteins in different organisms have different distributions of radii of gyration *R_g_*. We perform two-sample Kolmogorov–Smirnov (KS) test to compare the *R_g_* distributions of the proteins in different organisms (details listed in SI Methods). It is observed that some organisms have very similar *R_g_* distributions, and there are no statistically significant differences between them (e.g., *M. jannaschii* vs. *E. coli*, and mouse vs. rat). However, for some distinct organisms, for example, prokaryotes vs. eukaryotes (e.g., *E. coli* vs. yeast), unicellular vs. multicellular organisms (e.g., yeast vs. mouse), or multicellular organisms with significantly different numbers of cell types (e.g., *C. elegans* vs. human), there are statistically significant differences between them. More interestingly, if we sort the organisms according to the median *R_g_*, it is observed that larger median *R_g_* correlate with increasing organismal complexity. To evaluate such a correlation, for a given organism proteome, we introduce the total number of proteins and the total chain length of the proteins as measures of organismal complexity. As shown in Fig. 1B, both complexity measures are proportional to the median *R_g_* for proteins with similar chain lengths, demonstrating the correlation between protein flexibility and organismal complexity. Note that such a correlation is robust. For example, if one considers the proteins with other given chain lengths or selects other structural descriptors such as the solvent-accessible surface areas (SASA) to quantify the flexibility of the proteins, similar correlations can also be observed (SI Fig. S2).

**Figure 1.**
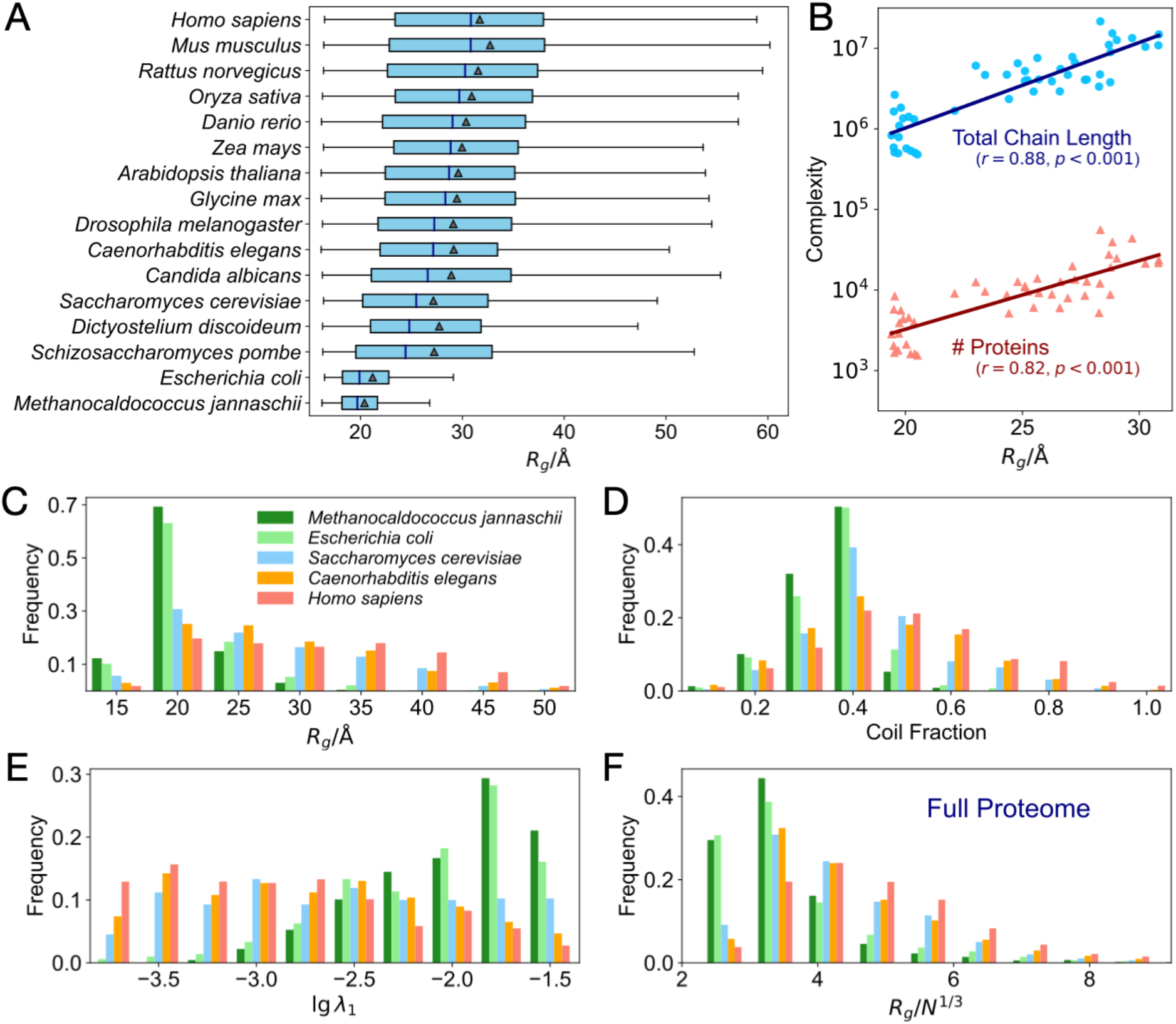
The statistics of AlphaFold-predicted structures indicate the correlation between the flexibility of constituent proteins and the complexity of organisms. (A) For the proteins from 16 model organisms with similar chain lengths *N* ≈ 250 (225 *≤N* < 275), the distributions of the radii of gyration *R_g_* are shown as the box-and-whiskers (extreme values not shown). Here, the triangle and the vertical bar in the box denote the mean value and the median of the *R_g_*, respectively. (B) For all the 48 organisms in AlphaFold DB, the measures of organismal complexity (the total number of proteins and the total chain length of the proteins in the organism proteome) vs. median *R_g_* of the proteins with chain lengths *N* ≈ 250. For proteins from the five selected organisms with chain length *N* ≈ 250, the histogram of the (C) radii of gyration *R_g_*, (D) coil fraction, and (E) the slowest mode eigenvalue at the logarithmic scale. (F) The distribution of the normalized *R_g_* (i.e., *R_g_/N^1/3^*) for all the proteins in the proteome of the five selected organisms.

To demonstrate such a correlation clearly, we select five species with different organismal complexity for detailed examinations. The five selected species include archaea (*M. jannaschii*), bacteria (*E. coli*), unicellular eukaryotes (budding yeast, *S. cerevisiae*), multicellular invertebrates (*C. elegans*), and human (*Homo sapiens*) Based on the proteomes of these organisms, we select the proteins with chain lengths *N ≈* 250, conduct structural analysis, and perform comparative analysis. The histograms shown in Figs. 1C-E demonstrate how the shape, secondary structures, and equilibrium dynamics of the proteins vary in the selected organisms.

First, as shown in Fig. 1C, for the proteins with similar chain lengths, the mean value, as well as the standard deviation of the radius of gyration *R_g_* gradually increases for organisms with higher complexity. Then, we apply the DSSP algorithm (Kabsch and Sander 1983; Joosten et al. 2011) to assign secondary structures (helices, sheets, or coils) to the proteins and investigate how the structural flexibility of the constituent proteins varies in different organisms. We find that the average fraction of coils increases as the organismal complexity increases (Fig. 1D). Next, we examine the dynamical flexibility of the proteins according to their vibrations around the native structure. These vibrations are closely related to the functional dynamics of the proteins, and can be predicted by the elastic network model (ENM) (Haliloglu et al. 1997; Bahar et al. 1998; Bahar et al. 2010). Based on ENM, one can conduct the normal mode analysis and obtain the eigenvalues corresponding to the vibration modes (Case 1994; Hayward et al. 1995; Wako and Endo 2017). Among these eigenvalues, the smallest nonzero eigenvalue *λ*_1_, which corresponds to the square of the slowest mode frequency 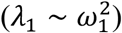 and is proportional to the inverse square of the amplitude of the motion 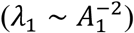, can be recognized as a measure of dynamical flexibility. As shown in Fig. 1E, statistically, for organisms with higher complexity, their constituent proteins will exhibit slower vibrations (i.e., smaller *Λ*_1_) around the equilibrium. Usually, the slow vibrations are closely related to the high modularity of the residue contact networks. We also show that for the proteins with similar chain lengths, the mean value and the standard deviation of modularity increase as the organismal complexity increases (see SI Figs. S3 and S4). In short, the above results clearly demonstrate the correlation between the flexibility of constituent proteins and the complexity of organisms. Further analyses show that even if we remove the proteins with low prediction confidence from the dataset, a similar correlation can still be observed (see Discussions and SI Fig. S5).

Furthermore, similar structural statistics can be extended to the analysis of the full proteomes. To compare the proteins with different chain lengths, we normalize the radii of gyration as *R_g_*/*N*^1/3^. When a protein is highly ordered and densely packed into a globular shape, then the normalized *R_g_* will be small; in contrast, when a protein is highly disordered or deviates from a globular shape, then the normalized *R_g_* will be large. The statistics of the normalized *R_g_* are shown in Fig. 1F. For the five selected organisms, the statistics of the full proteome show a similar trend as the statistics of the proteins with comparable chain lengths. For organisms with higher complexity, there will be larger mean values and standard deviations of the normalized *R_g_*. In SI Fig. S2, we conduct similar statistics for the full proteomes of all the 48 organisms in AlphaFold DB, compare the differences between the normalized *R_g_* distribution of different organisms, and show the correlations between the normalized *R_g_* and the complexity measures. These results further suggest that the increasing organismal complexity is accompanied by higher flexibility of the proteins in the full proteome. Further analyses (see SI Fig. S4) also show that structural diversity (measured by the standard deviation of the normalized *R_g_*) of constituent proteins also correlates with organismal complexity.

### The scaling analyses suggest a decreasing fractal dimension of constituent proteins as the organismal complexity increases

Based on the Protein Data Bank (PDB) (Berman et al. 2000), recent scaling analyses (Tang et al. 2017; Tang and Kaneko 2020) revealed the power laws related to the size dependence of the protein structure and dynamics. Due to the limited number of experimentally determined protein structures, it had been unable to compare the scaling coefficients for proteomes of different organisms separately. AlphaFold DB, however, compensates for the lack of experimental data, enables us to perform accurate scaling analyses of proteins across different organisms, compare scaling coefficients, and reveal the correlation between the organism’s complexity and the structural dynamics of their constituent proteins.

In this work, based on the predicted structures from AlphaFold DB, we apply the scaling analyses to the proteins from different species. For proteins within an organism, we first divide the proteins into bins according to their chain lengths. Then, for the proteins in the same bin (i.e., with similar chain length *N*), we calculate the mean value, as well as standard error, of the length of the shortest principal semi-axis *L_c_* and the slowest mode eigenvalue *λ*_1_. In this way, we obtain the size dependence of proteins’ shape and dynamics. Figure 2A and B show the results of the scaling analysis for the five selected organisms.

**Figure 2.**
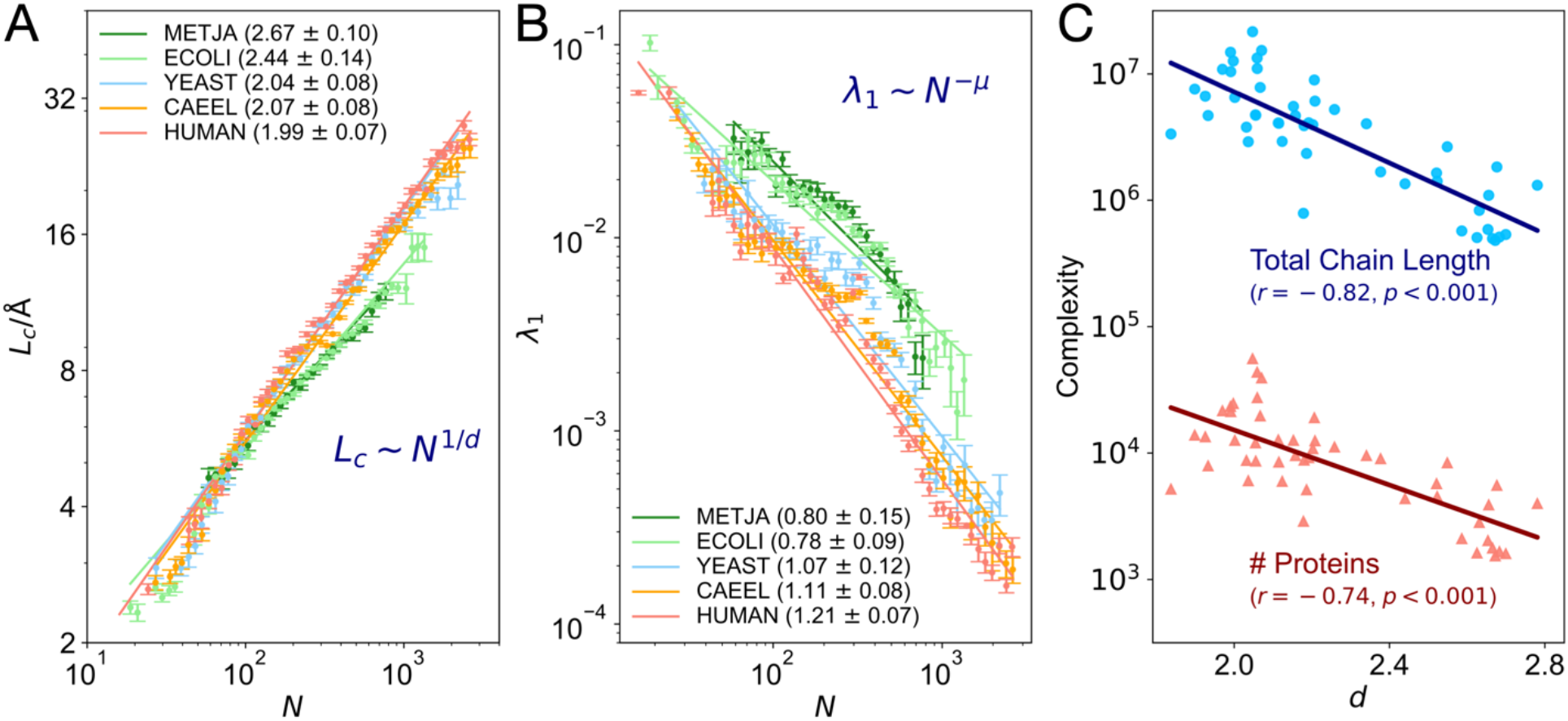
The scaling analyses suggest a decreasing fractal dimension of constituent proteins as the organismal complexity increases. For the proteins from five selected organisms, the size (chain length *N*) dependence of (A) length of the shortest semi-axis *L_c_* and (B) the slowest mode eigenvalue *λ_±_*. Here, for the data in all the bins, the average standard error of *L_c_* is below 3.5% of the mean value, and the average standard error of is below 13.1% of the mean value. The fitted scaling coefficients *d* (fractal dimension) and *μ* are listed in the legends of the subplots (A) and (B), respectively. These scaling coefficients are obtained by robust fittings with Theil–Sen estimators (95% confidence interval). (C) For all the 48 organisms in AlphaFold DB, the measures of complexity (the total number of proteins and the total chain length of the proteins in the organism proteome) vs. the estimated fractal dimension *d*.

Let us first analyze the scaling relations between protein shape and chain length *N* for different species. As the protein shape is described by the length of the shortest principal semi-axis *L_c_*, there will be a scaling relation: *L_c_* ~ *N^1/d^*, where *d* is the average fractal dimension of the proteins. For example, when a protein is densely packed into a globular shape in three-dimensional space, there will be *L_c_* ~ *R_g_* ~ *N*^1/3^ (i.e., *d* = 3). The scaling analyses shown in Fig. 2A indicate that the average fractal dimensions *d* vary across different organisms. Besides, for all the organisms, it is observed that *d* ≤ 3, displaying that the proteins are not always folded into densely packed globules. Previous studies on the fractal structure of the proteins (Lewis and Rees 1985; Liang and Dill 2001; Reuveni et al. 2008), as well as the statistics of the normalized *R_g_* (see Fig. 1F), can support such a result.

Then, we investigate the size dependence of the equilibrium dynamics of the native proteins. Previous research based on the PDB had shown that as the protein chain length *N* increases, the vibrations of the protein would become slower (Tang and Kaneko 2020), i.e., the slowest mode eigenvalue versus chain length *N* obeys the scaling relation: ~ *N*^−*μ*^, where *μ* ≈ 1. Notably, based on the AlphaFold DB, as shown in Fig. 2B, detailed analyses reveal that the scaling coefficients *μ* vary across different organisms. Next, let us take a closer look at the scaling coefficients *d* in Fig. 2A, one can find that, for two organisms with significant complexity differences (e.g., *E. coli* vs. human), there are significant differences in the corresponding fractal dimension *d*. The average fractal dimension of the constituent proteins of human is significantly lower than that of *E. coli*. According to such a result, one may conjecture that the average fractal dimension of constituent proteins is negatively correlated with the organismal complexity. To validate such a conjecture, we estimate the average fractal dimension *d* for all the 48 species in AlphaFold DB and evaluate their correlation with the complexity measures (total kinds of proteins and total chain lengths of proteins) of the organisms. The fittings in Fig. 2C confirm such a conjecture. The negative Pearson correlation coefficients indicate that for the species with higher organismal complexity, their constituent proteins will be lower average fractal dimension *d*. Such a correlation can also be validated by the statistics of the fractal dimension (i.e., the average packing dimension calculated with the boxcounting method) for all the proteins (see SI Fig. S1C). Besides, for a given proteome, the average fractal dimension of the proteins can also be estimated by the size dependence of the modularity *Q* of the proteins’ residue contact networks (Guimerà et al. 2004; Tang and Kaneko 2020). Although the average fractal dimension 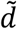 (estimated from *N* vs. *Q*) may be different from *d* (estimated from *N* vs. *L_c_*), it is also observed that a decreasing fractal dimension 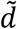 correlates with the increase of organismal complexity (see SI Fig. S3B). Similarly, we evaluate the correlation between the scaling coefficient *μ* and organismal complexity. As shown in SI Fig. S3D, the scaling coefficient *μ* is correlated with the measures of organismal complexity, suggesting that the constituent proteins of organisms with higher complexity will have slower vibration frequencies.

### Higher organismal complexity correlates with higher functional specialization and lower dimensionality in protein dynamics

According to the “structure-dynamics-function” paradigm (Friedman 1985; Agarwal 2006; Haliloglu and Bahar 2015), the dynamics encoded by the three-dimensional structure determine the functionality of the protein. As reported by previous research, for proteins with ordered native structures, the dynamics are suggested to be constrained to low-dimensional manifolds and could be described by a few ‘slow modes’ which correspond to low vibration frequency, low excitation energies, and large amplitudes motions. These slow modes are robustly encoded in the native structure of proteins. One may conjecture that the slow mode distributions of the constituent proteins are also correlated with the organismal complexity. To validate such a conjecture, for every protein, we construct the corresponding elastic network and conduct the normal mode analysis. In this way, the vibration spectrum can be obtained (see Method). In the spectrum, the eigengap between the leading two eigenvalues (*λ*_2_ and *λ*_1_) reflects how much the equilibrium dynamics of the protein will be dominated by the slowest normal mode (Togashi et al. 2007). For example, if *λ*_2_/*λ*_1_ = 10, then the ratio of the square amplitudes of the two modes will be: 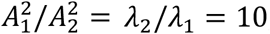, that is, the motion along the direction of the first mode is an order of magnitude greater than the motion along the direction of the second mode, reflecting a high functional specialization of the protein, or say, low dimensionality in protein dynamics. In contrast, if *λ*_2_/*λ*_1_ ≈ 1.01, the protein has almost the same tendency to move along the directions associated with the first and second slowest modes, indicating a higher dimensionality in dynamics and a lower functional specialization of the proteins. In short, the eigengaps between the leading eigenvalues and dimensionality in protein dynamics can act as measures of the functional specialization of the proteins.

Similar to previous sections, for the proteins with similar chain lengths *N* ≈ 250 from the five selected organisms, we conduct normal mode analysis based on the elastic network model and calculate the corresponding eigenvalues. Then, the distributions of the eigengaps *λ*_2_/*λ*_1_, *Λ*_3_/*λ*_2_, and *λ*_4_/*λ*_3_ (at logarithmic scale) are obtained. As shown in Figs. 3A-C, all the average ratios between neighboring eigenvalues increase as the organismal complexity increases, clearly demonstrating the increasing scale separation in the vibrational frequencies of the constituent proteins. This result indicates that, for organisms with higher complexity, their constituent proteins will have higher functional specialization and lower dimensionality. Moreover, it is worth noting that similar correlations can happen on every scale. As the organismal complexity increases, the constituent proteins have a statistical trend that the motions along the direction of the first modes have evolved to be much greater than the second, the motions along the second modes have evolved to be much greater than the third, etc.

**Figure 3.**
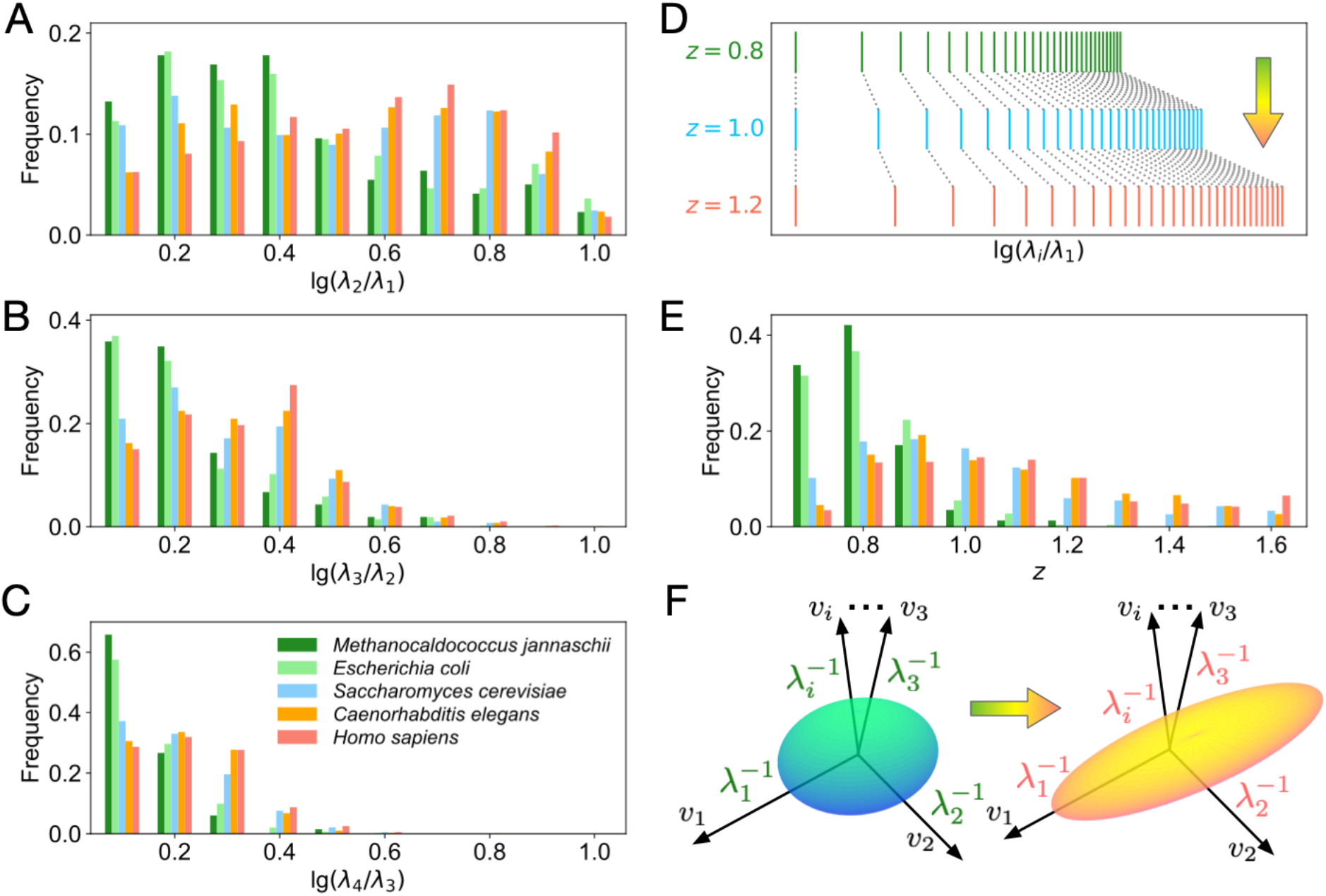
The increasing eigengaps in the vibration spectra imply that higher organismal complexity correlates with higher functional specialization and lower dimensionality in protein dynamics. For proteins with similar chain lengths *N* (225 ≤*N* < 275) in the five selected organisms, the distribution of the eigengaps (A) *λ*_2_/*λ*_1_, (B), *λ*_3_/*λ*_2_ and (C) *λ*_4_/*λ*_3_ at the logarithmic scale. (D) Illustration of power-law spectra at the logarithmic scale with Zipf’s coefficients *z* = 0.8, 1.0, and 1.2. (E) For proteins with similar chain lengths in different species, the distribution of the Zipf’s coefficients *z*. (F) Illustration of the trend of dimensional reduction in the conformational spaces. The arrows in subplots (D) and (F) indicate the increase in organismal complexity.

To quantify such a scale-free property, it is necessary to introduce the power laws to describe the vibrations of the proteins. Previous studies have shown that the vibrational spectrum of proteins obeys a power-law distribution (Reuveni et al. 2008), and the rank-size distribution of the inverse of eigenvalues follows a Zipf-like distribution (Tang et al. 2020; Tang and Kaneko 2021; Xie et al. 2022). In SI Fig. S6, several proteins are provided as examples. These proteins have similar chain lengths, but their vibration spectra are different. The rank-size distribution of the slow-mode eigenvalues shows a power-law-like behavior, which can be described as the Zipf’s law, i.e., 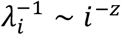, where *z* is known as the Zipf’s coefficient.

Generally, for the proteins with similar sizes, a larger *z* corresponds with larger eigengaps (Fig. 3D). In Fig. 3E, the statistics of Zipf’s coefficients *z* show that, for organisms with higher complexity, *z* will be larger, i.e., there will be larger eigengaps in the vibrational spectra of constituent proteins. Such a correlation is illustrated in Fig. 3F. In the illustration, the conformational space (the space encompassing all possible structures results from thermal fluctuations) of a protein around its native structure is represented as an ellipsoid. The axes of the ellipsoids denote the normal modes, and their semi-length represents the square magnitudes of the motions in the directions of the normal modes. The functional specialization and low dimensionality correspond to the anisotropy of the conformational space. As the organismal complexity increases, both the structure and dynamics of the constituent proteins show a statistical trend towards dimensional reduction.

### Organismal complexity correlates with the assortativity of the residue contact networks and the hydrophilic-hydrophobic segregation in protein sequences

In previous studies, residue contact networks of native proteins have been shown to play a dominating role in determining protein dynamics and functions (Bahar et al. 2010; Atilgan et al. 2012). The residue contacts are, in turn, largely determined by the protein sequences. Therefore, we may use the residue contact network as a bridge to investigate how organismal complexity correlates with the sequences of the constituent proteins.

In the previous subsection, we show that higher organismal complexity correlates with larger scale separation in the vibrational frequencies of the proteins. In fact, such a correlation in frequency space can correspond to the changes in the distribution of the residues’ local packing density in the real space. According to the local density model (Halle 2002), there is a nearly linear relationship between the residues’ square of vibration frequency and the local packing density. As a result, the increasing eigengaps in vibrational frequencies corresponds to the increasing variances of the residues local packing density. In protein molecules, the residue packing density is mainly determined by the hydrophobic effects. As illustrated in Fig. 4A, by avoiding exposure to water, hydrophobic residues are buried in the interior of the protein, forming the hydrophobic cores with high local packing density. In contrast, the hydrophilic (polar) residues are likely to exposure to the solvent and have low local packing density. It is such hydrophobic effects that serve as the major driving force of protein folding (Dill 1990; Dill and MacCallum 2012).

**Figure 4.**
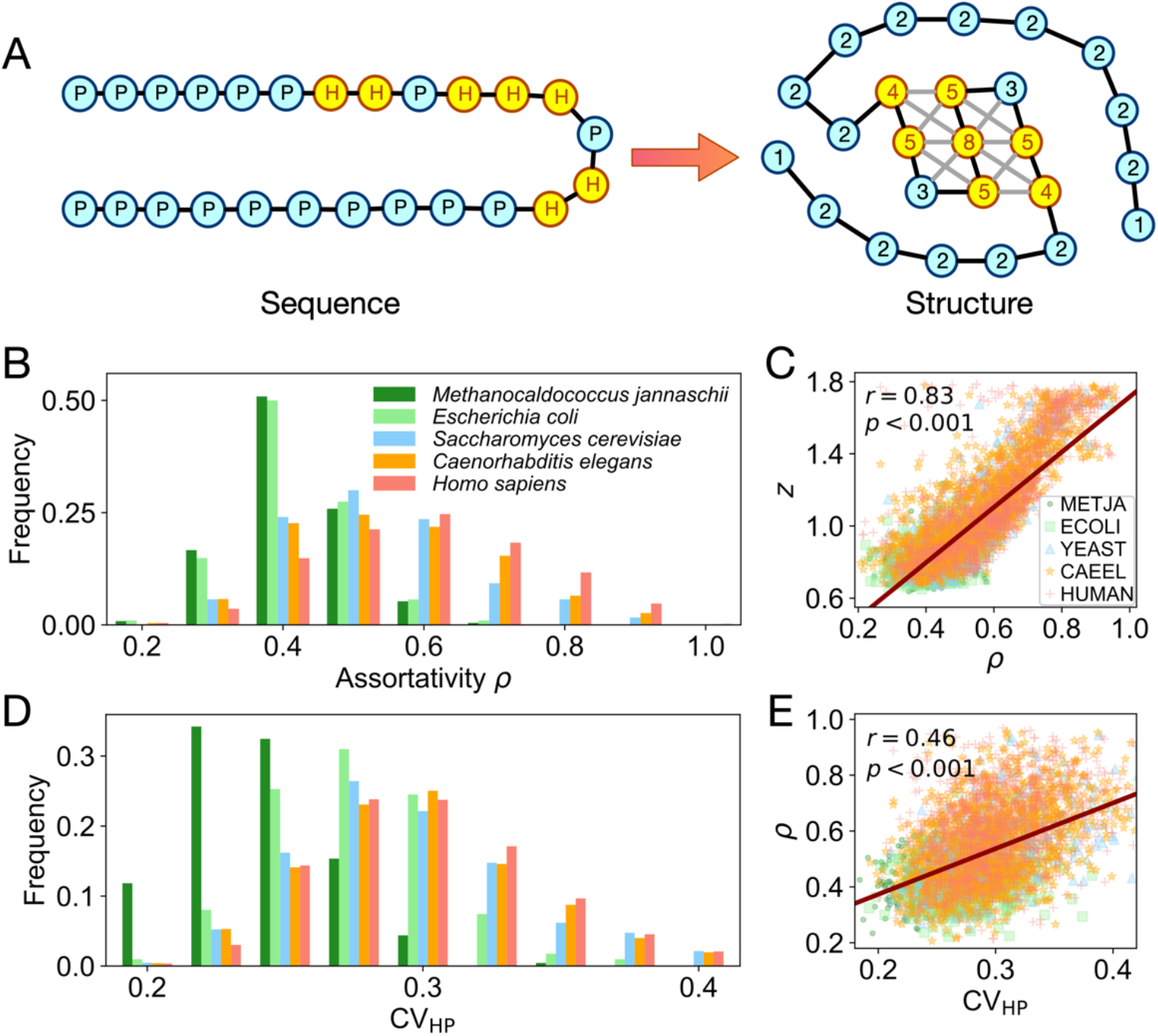
Organismal complexity correlates with the assortativity of the residue contact networks and the hydrophilic-hydrophobic segregation in protein sequences. (A) Illustration of a protein sequence (left) folds into a native structure described as an assortative residue contact network (right). The yellow and cyan nodes denote the hydrophobic (H) and polar (P) amino acid residues, respectively. When the protein folds into the native structure, the hydrophobic residues tend to aggregate into a densely connected hydrophobic core. In the subplot (right), the numbers represent the node’s degree (number of connections with other nodes). (B) For proteins with similar chain length *N* (225 ≤*N* < 275) in the five selected organisms, the histogram of residue contact network assortativity *ρ*. (C) The scattering plot and the fitted trend line of the assortativity *ρ* vs. Zipf’s coefficient *z*. (D) For proteins with similar chain lengths *N* in different organisms, the histogram of the hydropathy variations CV_HP_. (E) The scattering plot and the fitted trend line of hydropathy variation CV_HP_ vs. residue contact network assortativity *ρ*.

For the proteins with similar chain lengths, the variance of the local packing density is closely related to the assortativity *ρ* of the residue contact network. The assortativity of a network is defined as the Pearson correlation coefficient of degrees between pairs of linked nodes (Newman 2002; Newman 2003). According to the definition, if the residues (nodes) with high packing density (degrees) are more likely to have contact with each other, then there will be a higher assortativity *ρ* and a larger variance in the packing density distribution of residues. Remarkably, as shown in Fig. 4B, as the organismal complexity increases, the constituent proteins with similar chain lengths will have a larger mean value of assortativity *ρ*. Notably, such a correlation is in line with the trend related to the increasing eigengap of vibration spectra. As shown in Fig. 4C, for the proteins in our dataset, regardless of the species they belong to, the assortativity *ρ* is proportional to Zipf’s coefficient *z*. This result indicates that we have found the topological descriptor associated with the functional specialization of the protein.

Next, let us discuss how organismal complexity correlates with the sequences of the constituent proteins. As illustrated in Fig. 4A, the spatial segregation of hydrophilic and hydrophobic residues is closely related to their segregation in sequence. When hydrophilic and hydrophobic residues are uniformly mixed and randomly distributed in the sequence, they will be less likely to have profound spatial segregation. Conversely, when there are longer hydrophilic or hydrophobic fragments in a sequence, the hydrophobic cores can be formed more easily. Here, we introduce the hydropathy variation CV_HP_ to quantify the hydrophilic-hydrophobic segregation in protein sequences, where CV_HP_ is defined as the coefficient of variation (i.e., standard deviation divided by the mean value) of the filtered hydropathy profile of a sequence (see Method). As shown in Fig. 4D, for proteins with similar chain lengths, as the complexity of the organism increase, there will be a larger average CV_HP_, which corresponds to a more significant hydrophilic-hydrophobic segregation. Further evaluation of such a correlation is shown in SI Fig. S8. As shown in Fig. 4E, for proteins in our dataset, regardless of the organisms they belong to, the hydropathy variation CV_HP_ is proportional to the assortativity of the residue contact network, implying that the spatial segregations of hydrophilic and hydrophobic residues do have statistically significant correlation with the sequence segregations.

The sequence analysis result also uncovers the differences between the constituent proteins from archaea (*M. jannaschii*) and bacteria (*E. coli*). Although their structures (e.g., radii of gyration) do not show significant differences (see Fig. 1C), their sequences do show significant differences (Fig. 4D). These results do not mean that *E. coli* has a significantly higher complexity than *M. jannaschii*. It is their living environments that determine the major difference in sequence. As a thermophilic methanogenic archaean, *M. jannaschii* lives in extreme environments such as hypothermal vents at the bottom of the oceans. The optimal growth temperature of *M. jannaschii* is about 85°C (Jones et al. 1983; Bult et al. 1996), which is even higher than the melting temperature (~50°C) of most proteins in *E. coli* (Ghosh and Dill 2010). The low hydropathy segregation in the protein sequences of *M. jannaschii* contributes to the stabilization of the surface residues of the proteins, thus leading to the organism’s adaptation to a hot environment.

### Phylogenetic proximity of homologous proteins correlates with the statistical structural proximity of the proteomes

In previous sections, our statistical structural analysis of proteomes has shown that the organismal complexity can be reflected by the structural properties of the constituent proteins. Since the complexity of organisms has emerged through evolution, one may conjecture that the phylogenetic proximity can be reflected in the structural statistics of the constituent proteins. To validate such a conjecture, here we define the statistical structural proximity between two proteomes based on the Kolmogorov-Smirnov (KS) statistic *D* (Massey 1951) between the distributions of the radii of gyration *R_g_* for proteins with similar chain length *N* ≈ 250. Intuitively, the KS statistic *D* can be understood as the largest absolute difference between two cumulative distribution functions. When two proteomes have very similar *R_g_* distributions, then their distance (KS statistic *D*) will be small, and the statistical structural proximity between the two proteomes will be high. By calculating the KS statistic between every two organisms, a statistical structural distance matrix can be constructed. Interestingly, the phylogenetic tree (Fig. 5A) generated with the sequences of a homological gene shows correspondence with the statistical structural distance matrix. The lineages that are close in the phylogenetic tree also have a small distance (i.e., small KS statistic *D*) between the proteomes. Such a result not only demonstrates that phylogenetic proximity correlates with the statistical structural proximity of the proteomes, but also suggests that there is a statistical trend of protein evolution in parallel with organism evolution.

**Figure 5.**
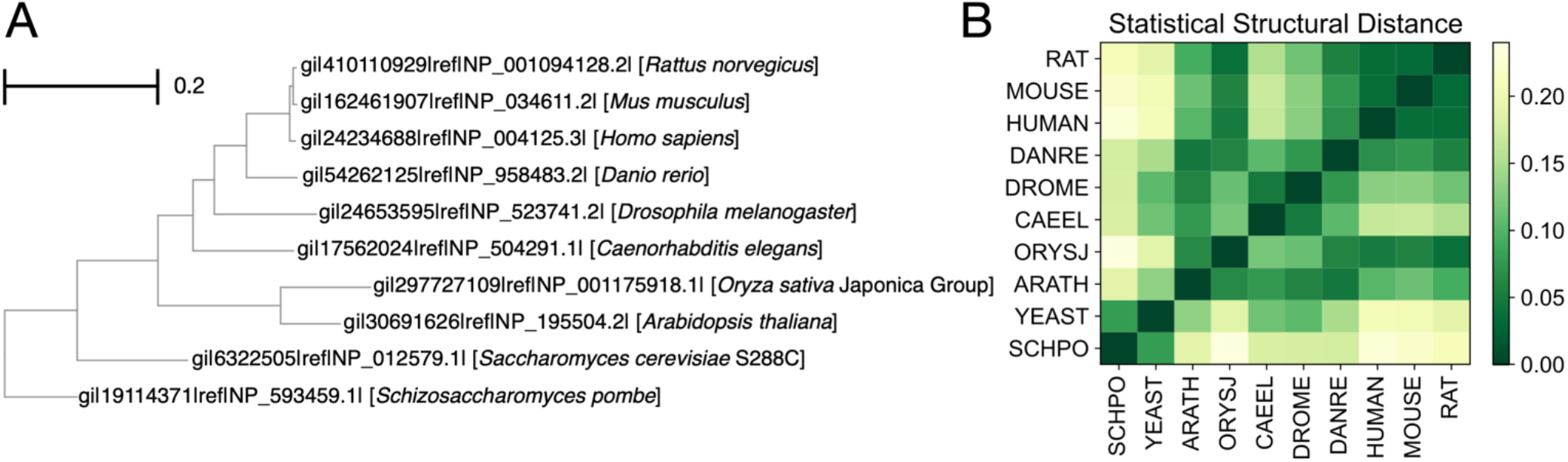
Phylogenetic proximity of homologous proteins correlates with the statistical structural proximity of the proteomes. (A) A phylogenetic tree generated with the sequences of the heat shock protein (HSPA9, HomoloGene: 39452) by NCBI protein BLAST (Gish and States 1993; McGinnis and Madden 2004). (B) The statistical structural distance matrix with entries of the KS statistic *D* between *R_g_* distributions of the organisms’ constituent proteins with similar chain length *N* ≈ 250.

Notably, the matrix shown in Fig. 5B is based on a simplistic definition of statistical structural proximity between proteomes, say, only *R_g_* for proteins with similar chain lengths are considered. One may take into account proteins with other chain lengths or introduce other structural descriptors (e.g., SASA, secondary structures, modularity etc.) to refine the definition. Further analysis (see SI Fig. S9) shows that, even though statistical structural proximity based on other descriptors may vary in values, we can generally observe that the lineages that are close in the phylogenetic tree show high statistical structural proximity. In all, the statistical structural proximity across the proteomes may indirectly reflect the evolutionary relationships among species.

## Discussion

In this work, based on the protein structures predicted by AlphaFold, we propose a statistical framework for proteome analysis by comparing the protein structures in different organisms. Rather than studying the evolution of specific protein families or superfamilies, this study aims to reveal the correlations between organismal complexity and structural or dynamical properties of the constituent proteins. It is observed that, statistically, the constituent proteins of higher-complexity organisms will have higher flexibility, lower fractal dimensions in both structure and dynamics, and higher degrees of hydrophobicity segregation in both their structures and sequences. Note that these correlations do not depend on the definition of complexity. Although the mathematical definition of complexity is still controversial (Lloyd 2001), different measures of organismal complexity (e.g., number of cell types, genome size, proteome size, etc.) are correlated (Markov et al. 2010; Niklas et al. 2014). Besides, it is worth noting that the structure prediction confidence of AlphaFold will not affect our main conclusions. For the AlphaFold-predicted structures, lower prediction confidence (which can be quantified as lower pLDDT values, see SI) usually arrives for proteins with long disordered regions (Jumper et al. 2021). Interestingly, even if we remove those proteins with lower prediction confidence (i.e., with long intrinsic disorder regions) from the dataset, similar correlations can still be observed (see Fig. S2). Moreover, our study focuses primarily on the global dynamics (e.g., slowest modes) that are robust to the local variations in native structures (Bahar et al. 2010; Tang et al. 2020). Such insensitivity to prediction confidence can further strengthen our conclusions.

Our topological and sequence analyses show that higher organismal complexity correlates with the assortativity of the residue contact networks and the hydrophilic-hydrophobic segregation in protein sequences. These correlations are consistent with what has been suggested by previous evolutionary studies (Phillips 2009; Phillips 2012; Hemery and Rivoire 2015; Foy et al. 2019; Moret et al. 2019; Phillips 2020), and may guide the design or modification of proteins. For example, one may enhance the flexibility or functional sensitivity of a protein by greater segregation of hydrophobic and hydrophilic amino acids in the sequence. Besides, hydrophilic-hydrophobic segregation can also prevent the dysfunctional aggregation of proteins (Foy et al. 2019). Notably, the sequence analysis does not rely on AlphaFold predictions in any way, which acts as essential support that our main conclusions do not arise from a systematic bias inherent in structural prediction methods, but reflect the natural tendency.

Our analysis also shows that the phylogenetic proximity of homologous proteins correlates with the statistical structural proximity of the proteomes, indicating the evolutionary roots of the statistical trend discussed in this paper. Although not all the constituent proteins in an organism will evolve in the same direction, statistically, as the organismal complexity increases, the constituent proteins show higher flexibility. The evolution of many protein families is in line with this statistical trend (Brocchieri and Karlin 2005; Kasho et al. 2006). Besides, such a trend is also consistent with the fact that the intrinsic disorder is more abundant in organisms with higher complexity (Niklas et al. 2014; Basile et al. 2019). Not only will the intrinsic disorder proteins (or regions) exhibit high dynamical plasticity, as shown in previous research (Meier and Özbek 2007; Tokuriki et al. 2008; Tokuriki and Tawfik 2009; Marsh and Teichmann 2014), but they may also exhibit high evolvability (towards more specialized function and towards new folds). Still, it is worth noting that, as thermostability becomes a selection pressure, the proteins can evolve to be less flexible (Berezovsky and Shakhnovich 2005). This statistical trend is also parallel with the phenomenon of evolutionary dimensional reduction observed in other experimental and theoretical studies (Hemery and Rivoire 2015; Dutta et al. 2018; Furusawa and Kaneko 2018; Eckmann et al. 2019; Sakata and Kaneko 2020; Sato and Kaneko 2020; Eckmann and Tlusty 2021). In protein dynamics, it is also implied in the emergence of the funnel-like energy landscape (Onuchic et al. 1997), and may contribute to the efficiency and robustness of the protein function. For allosteric proteins, it is observed that evolution designs the sequence and shapes the structure of the proteins, leading toward more specific transition pathways (Togashi et al. 2007; Li et al. 2011).

Moreover, the statistical correlation between proteins’ functional specialization and organismal complexity is in line with the experimental observations that ancestral enzymes are likely to have high promiscuity, as they may be able to catalyze a wide variety of chemical reactions (O’Loughlin et al. 2006; Khersonsky and Tawfik 2010; Takano et al. 2013; Petrovic et al. 2018; Gardner et al. 2020; Modi et al. 2021). It is widely observed that ancient generalist proteins tend to evolve toward specialists (Soskine and Tawfik 2010). Previous research in designing thermally stable and promiscuous enzymes with ancestral sequence reconstruction can act as additional support of such a trend (Wheeler et al. 2016; Trudeau and Tawfik 2019; Pinto et al. 2022). Notably, the correlation between protein flexibility and functional specialization is also statistical. There are counterexamples that proteins with higher conformational plasticity show higher promiscuity (Campbell et al. 2018).

The high thermal stability (which is usually associated with low flexibility and high promiscuity) of ancestral enzymes may in turn be compatible with the low complexity of the ancestral species. Organisms with lower complexity have relatively smaller genomes and fewer kinds of enzymes (Gagler et al. 2022). Despite a small genome size, promiscuous enzymes help these organisms achieve a variety of life activities. Conversely, larger genomes can encode more proteins capable of performing highly specialized functions and coping with more complex and diverse cellular environments. The specialization and diversification of proteins enable them to function in a more complex and diverse cellular environment. Consequently, complex organisms can perform their biological functions more effectively, acquiring the plasticity to adapt to complex and diverse external environments. The compatibility between organismal complexity and functional specialization of constituent proteins suggests the interdependence between the whole and parts for biological systems and other kinds of complex systems, shedding light on the design of complex systems. As a system becomes more complex, its components or elements should change their properties (e.g., become more plastic or modularized).

In summary, based on the AlphaFold DB, we establish a framework for the comparative analyses of the constituent proteins of different organisms. The statistics of AlphaFold-predicted structures have revealed the correlations between the complexity of organisms and features related to the structures (topology), dynamics, and sequences of constituent proteins. Moreover, we show that the phylogenetic proximity is correlated with the statistical structural proximity of the organisms’ proteomes, indicating the connections between protein evolution and organism evolution. Our analysis suggests a statistical trend in protein evolution, that is, as organisms evolve toward higher complexity, their constituent proteins may evolve toward higher flexibility and structural diversity, statistically. In the future, the proteome analysis based on AI-predicted protein structures, integrated with other kinds of bioinformation such as protein-protein interactions (Maslov and Sneppen 2002; Zhang et al. 2008), expression levels (Drummond et al. 2005), evolutionary speed (Agozzino and Dill 2018), etc., will definitely offer us new insight into the behaviors and evolution of cells and organisms.

## Materials and Methods

### Data Availability

All the protein structures used in this study can be downloaded from AlphaFold Protein Structure Database (Varadi et al. 2022). A detailed description of the dataset is listed in SI Table S1. The curated data underlying this article are available in Github, at https://github.com/qianyuantang/stat-trend-protein-evo.

### Residue contact network

We construct the residue contact networks based on the native structure of the proteins. The residues (represented as their C_α_ atoms) are modeled as the nodes of the network. Our calculations of the modularity and assortativity are based on the residue contact networks corresponding to the native structure. When the mutual distance between two residues (nodes) is smaller than the cutoff distance *r_c_*, then the two residues will be connected with an edge. In this work, we take *r_c_* = 8 Å.

### Elastic network model (ENM)

By modeling the edges in the residue contact networks as linear springs, the elastic network models describe the equilibrium fluctuations of proteins as vibrations around the native conformation. These fluctuations are closely related to the functional dynamics of the proteins. In this work, our discussions are based on the Gaussian network model (GNM), the simplest form of the ENM, where the residue fluctuations are assumed to be Gaussian variables distributed around the equilibrium coordinates. The dynamics predicted by GNM can be well matched to experimental or simulation results (Haliloglu et al. 1997; Bahar et al. 2010). With GNM, the potential energy of a protein with chain length *N* is given as: 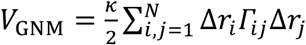, where *κ* is a uniform force constant; Δ*r_i_* and Δ*r_j_* denote the displacement of residues *i* and *j*, respectively; and *Γ_ij_* is the element of Kirchhoff matrix Γ. For residues *i* and *j* (*i* ≠ *j*), if their mutual distance *r_ij_* ≤ *r_c_*, then *Γ_ij_* = –1; if *r_ij_* > *r_c_*, then *Γ_ij_* = 0; and for the diagonal elements, *Γ_ii_* = – *k_i_* = — ∑_*j*≠*i*_ *Γ_ij_*, where *k_i_* denote the degree of node *i*. Here we take *r_c_* = 8 Å.

### Normal mode analysis

The eigendecomposition of matrix Γ gives the eigenvalues and the corresponding eigenvectors related to the motions of normal modes (Case 1994; Hayward et al. 1995; Wako and Endo 2017). To compare the eigenvalues for the proteins with different chain lengths, the diagonal elements of matrices Γ are normalized as 1. The normalized matrix is also known as the symmetric normalized graph Laplacian (Atilgan et al. 2012): *L* = *K*^-1/2^Γ*K*^−1/2^, where *K* is a diagonal matrix *K* = diag[*k*_1_, *k*_2_, – *k_N_*] describing the local packing density of the residues. Diagonalizing matrix *L*, we have *L* = *U*Λ*U^T^*, in which matrix 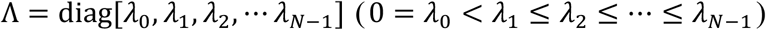 represents the eigenvalues, and matrix *U* = [*u*_0_, *u*_1_ *u*_2_, –, *u*_*N*−1_]^*T*^ denotes to the eigenvectors. The zeroth eigenvector *u*_0_ corresponds to the translational and rotational motions, other nonzero modes correspond to the vibrations of the proteins. The nonzero eigenvalue *λ_i_* is proportional to the square of the vibrational frequency *ω_i_*, i.e., 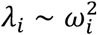. According to the equipartition of energy, the vibration amplitude *A_i_* is inversely proportional to the frequency: *A_i_* ~ 1/*ω_i_*. Thus, for two nonzero modes *i* and *j*, there is 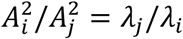. We use the Python package ProDy to calculate the normal modes of the proteins (Bakan et al. 2011).

### Power-law fitting and the Zipf’s coefficient

To fit the Zipf’s law 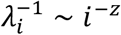 and obtain the coefficient *z*, in the calculation, we select the top 25% of the eigenvalues to perform the power-law fitting (Alstott et al. 2014). Note that the observed evolutionary trend as indicated by Zipf’s coefficient *z* are robust to the details related to the normal mode calculation and power law fitting (see SI Fig. S7).

### Hydropathy variation

In the calculation, we first obtain the original hydropathy data of a sequence according to the Zimmerman hydrophobicity scale of amino acid residues (Zimmerman et al. 1968). Then, a moving average with window length *l_w_* = 7 is introduced to calculate the hydropathy profile. If there are no significant hydrophobic-hydrophilic segregations in a sequence, the moving average will smooth out the differences between hydrophobic and hydrophilic amino acids, which will lead to little variation in the filtered hydropathy profile. In contrast, when there are significant hydropathy segregations in a sequence, there will be large variations in the filtered hydropathy profile. Thus, we introduce the coefficient of variation CV_HP_ (defined as the standard deviation divided by the mean value of the filtered hydropathy profile) to quantify the sequential segregation of hydrophobic and hydrophilic residues.

## Supporting information

Updated SI

## Acknowledgments

We gratefully thank Taro Toyoizumi, Xiangze Zeng, Hisao Moriya, Haobo Wang, Haiguang Liu, Zhengqi He, Sida Chen, Weiyi Qiu, Wenfei Li, Zeke Xie, Yingnan Li, Xinhong Liu, Lei-Han Tang and Xuefei Li for participating in stimulating discussions. This work was supported by Brain/MINDS from Japan Agency for Medical Research and Development (grant number JP21dm0207001), National Natural Science Foundation of China (grant number 11774157), a Grant-in-Aid for Scientific Research on Innovative Areas (grant number 17H06386) from the Ministry of Education, Culture, Sports, Science and Technology of Japan, a Grant-in-Aid for Scientific Research (grant number (A)20H00123) from the Japanese Society for the Promotion of Science, and Novo Nordisk Fonden (to K.K.).

