## Supplementary material for "The Statistical Trends of Protein Evolution: A Lesson from AlphaFold Database": Updated SI

### Methods

**Two-sample Kolmogorov–Smirnov (KS) test.** The two-sample KS test is a non-parameterized test that is usually applied to compare two samples and test whether the two samples come from the same distribution. In the KS test, the KS statistic $D$ is defined as the maximum difference (i.e., the vertical distance) between the two cumulative distribution functions (illustrated in Fig. S1B), and the $p$-value is a probability corresponding to $D$. The KS statistic $D$ can act as the measure of distance between two samples. For example, for proteins with similar chain lengths $N \approx 250(225 \leq N < 275)$, to compare the $R_g$ distribution of the proteins in two organisms A and B, we can take the protein $R_g$ data of organism A as sample A, and $R_g$ data of organism B as sample B. Then, in the two-sample KS test, a larger KS statistic $D$ with smaller $p$-value indicates more significant differences between the two samples. For the five selected organisms, the KS test results (KS statistics $D$ and $p$-values) of the structure- or sequence-related feature of the proteins from are listed in Table S2.

In testing whether two samples differ from each other, the null hypothesis (both samples come from a population with the same distribution) is rejected at level $\alpha$ when $D_{m,n,\alpha} > c(\alpha)\sqrt{\frac{n+m}{nm}}$, where $n$ and $m$ denote the size of the first and second sample, respectively, and $c(\alpha)$ denotes the critical value at level $\alpha$. In this work, we mainly consider the critical values $c(\alpha) = 1.628 (\alpha = 0.01)$, and $c(\alpha) = 1.949$ $(\alpha = 0.001)$. For the given two samples, we first calculate $\tau(\alpha) = \frac{D_{m,n,\alpha}}{c(\alpha)}\sqrt{\frac{nm}{n+m}}$, and if $\tau(\alpha) > 1$, then the null hypothesis can be rejected. For proteins with similar chain lengths $N \approx 250(225 \leq N < 275)$ from the 16 model organisms in the database, the results of the two-sample KS test (based on the distributions of $R_g$ and $R_g/N^{1/3}$) are shown in Fig. S10. A full version of the result, including the two-sample KS tests for 48 organisms in our database, are listed in the text files "KS-Rg-N250.txt" (based on the distribution of $R_g$) and "KS-Rg_norm-N_full.txt" (based on the distribution of $R_g/N^{1/3}$). In the text files, each row represents the KS test for the two samples of proteins from two organisms (species codes are listed). The corresponding KS statistics $D$, corresponding $p$-value, $\tau(\alpha = 0.01)$, and $\tau(\alpha = 0.001)$ are all listed in the file. The data files are available in Github, at https://github.com/qianyuantang/stat-trend-protein-evo.

**Modularity of the residue contact network.** Modularity is a topological descriptor that quantifies if a network can be easily divided into modules. It is defined as the fraction of the edges that fall within the given module minus the expected fraction when edges are distributed at random (Newman 2004; Newman 2006). According to this definition, for a network with $N$ nodes and $M$ edges described by the adjacency matrix $A$ (in which $A_{ij} = 1$ if and only if node $i$ and $j$ are connected), the modularity $Q$ can be calculated as $Q = Tr(x^T \cdot B \cdot x)/4M$, in which matrix $B$ is given as $B_{ij} = A_{ij} - k_i k_j/2M$, and vector $x$ is the column vector describing the network partition. For any given partition of a network, the corresponding modularity $Q$ can be calculated, and its value lies between $-1$ and 1. According to the definition, a network that can be more easily divided into modules would have a higher modularity $Q$. The appropriate partition of a network would maximize the modularity $Q$. In this work, we employ the Louvain method (Blondel et al. 2008) to partition the network and maximize the value modularity $Q$.

**Size dependence of modularity.** Usually, since lower-dimensional geometric graphs can be more easily divided into modules (e.g., a one-dimensional geometric graph can be divided into two modules by removing only one edge), networks embedded in a lower-dimensional space will have a higher $Q$. Theoretically, for a $d$-dimensional cubic lattice network with $N$ nodes, the modularity $Q$ vs. $N$ follows the scaling relation: $1 - Q \sim N^{-1/(\tilde{d}+1)}$, where $\tilde{d}$ can serve as another estimation of the average fractal dimension (Guimerà et al. 2004; Tang and Kaneko 2020). Note that even for the same group of proteins, the estimated $\tilde{d}$ may differ from the average fractal dimension $d$ (estimated from $N$ vs. $L_C$). Still, as shown in Fig. S3B, the estimated fractal dimension $\tilde{d}$ negatively correlates with the measures of organismal complexity, in line with the trend as shown in Fig. 2C of the Main Text.

**Confidence metric of structure prediction – pLDDT.** AlphaFold produces a per-residue confidence metric called the predicted local distance difference test (pLDDT) on a scale from 0 to 100. Usually, residues with pLDDT ≤ 50 correspond to very low confidence, residues with 50 < pLDDT ≤ 70 have low confidence, and residues with 70 < pLDDT ≤ 90 correspond to a generally correct backbone prediction, and residues with pLDDT > 90 are expected to be modeled to high accuracy. Although the residues with pLDDT < 50 are predicted with low confidence, low pLDDT values can work as reasonably strong predictors of disorder regions in the proteins, or say, those proteins with low average residue pLDDT usually have long disordered regions. In the Main Text, to show the correlation between organismal complexity and the flexibility of constituent proteins, our statistics also consider the AlphaFold-predicted structures with low average pLDDT values. It is worth noting that, even if we filter the proteins with low average pLDDT, as shown in Fig. S5, the statistical correlation is kept.

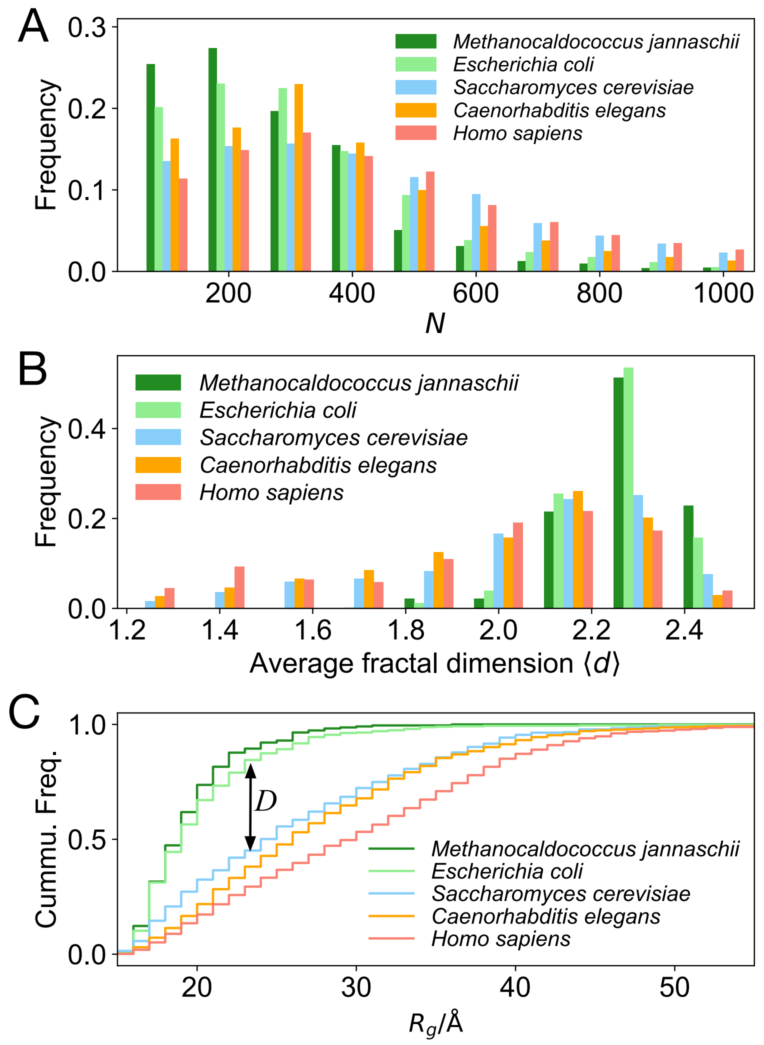


**Figure S1. Some more statistics of AlphaFold-predicted structures.** (A) For the five selected organisms, the statistics of the proteins' chain lengths. Note that the number of proteins with chain lengths $N$ around 200 to 300 is relatively high for all the selected species. Hence, in the main text, we mainly focus on the proteins with chain lengths $N \approx 250 (225 \leq N < 275)$. (B) For proteins from the five selected organisms with similar chain lengths $N \approx 250$, the histogram of the average fractal dimension $\langle d \rangle$. Here, to estimate the fractal dimension of a protein, we first calculate the fractal dimension of every residue based on the box-counting method (counting how the number of neighboring residues increases as the cutoff distance increases) and then calculate the average over all the residues. Note that there are many residues at the surface of the protein, so this estimation will give a relatively lower value of the fractal dimension. The result is consistent with the results shown in Fig. 2C of the Main Text. (C) For proteins from the five selected organisms with similar chain lengths $N \approx 250$, the cumulative distribution functions of the radii of gyration $R_g$. In this subplot, the distance $D$ (denoted by the two-way arrow) corresponds to the Kolmogorov–Smirnov (KS) statistics quantifying the distance between the distributions of the *E. coli*. and yeast (see the underlined entry in Table S2A).

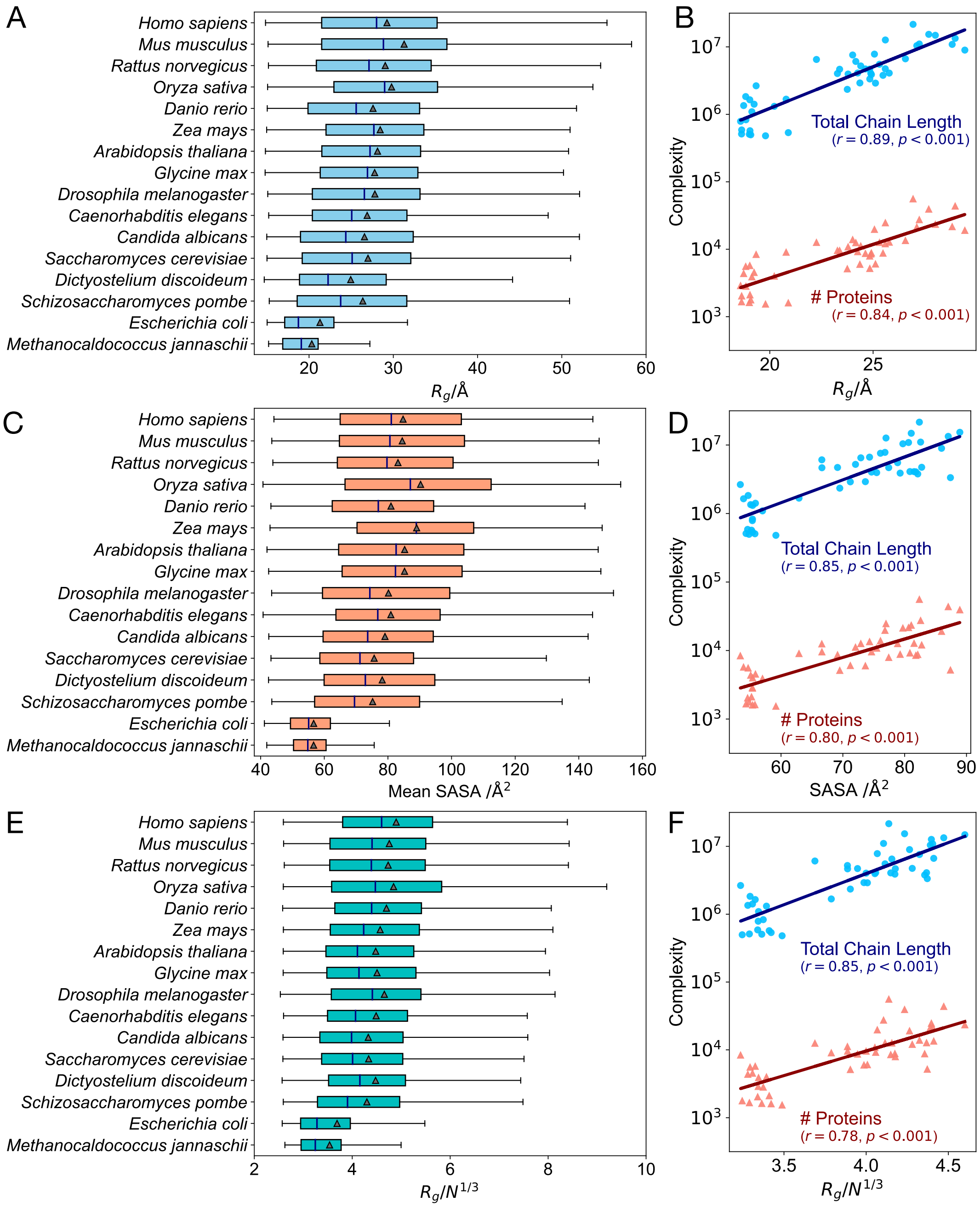


**Figure S2. The correlation between the flexibility of constituent proteins and organismal complexity is robust to the selection of chain length or indicators.** (A) For the 16 model organisms in AlphaFold DB, when selecting the proteins with some other chain lengths $N \approx 200$ ($175 \leq N < 225$), the distributions of the radii of gyration $R_g$ are shown as the box-and-whiskers (extreme values not shown). To compare with the results shown in the Main Text, here in all the subplots, the orderings of the species are based on Fig. 1A. The result indicates the correlation between the flexibility of constituent proteins and organismal complexity is robust to the selection of the protein sizes. (B) For the 48 organisms in AlphaFold DB, the measures of organismal complexity (the total number of proteins and the total chain length of the proteins in the organism proteome) vs. median $R_g$ of the proteins with chain lengths $N \approx 200$. (C) For the proteins with similar chain lengths $N \approx 250 (225 \leq N < 275)$ from 16 model organisms, the distributions of the solvent-accessible surface area (SASA) per residue are shown as the box-and-whiskers. (D) For the 48 organisms in AlphaFold DB, the measures of organismal complexity vs. median SASA of the proteins with chain lengths $N \approx 250$. These results in subplots (C-D) show that the observed correlation between the flexibility of constituent proteins and organismal complexity is robust to the selection of structure-related indicators. (E) For all the proteins from the proteome of the 16 model organisms, the distribution of the normalized $R_g$ (i.e., $R_g/N^{1/3}$) are shown as the box-and-whiskers. (F) For all the proteins from the proteomes of 48 organisms in AlphaFold DB, the measures of organismal complexity vs. median normalized $R_g$ of the proteins. The results suggest that the increasing organismal complexity is accompanied by higher flexibility of the proteins in the full proteome.

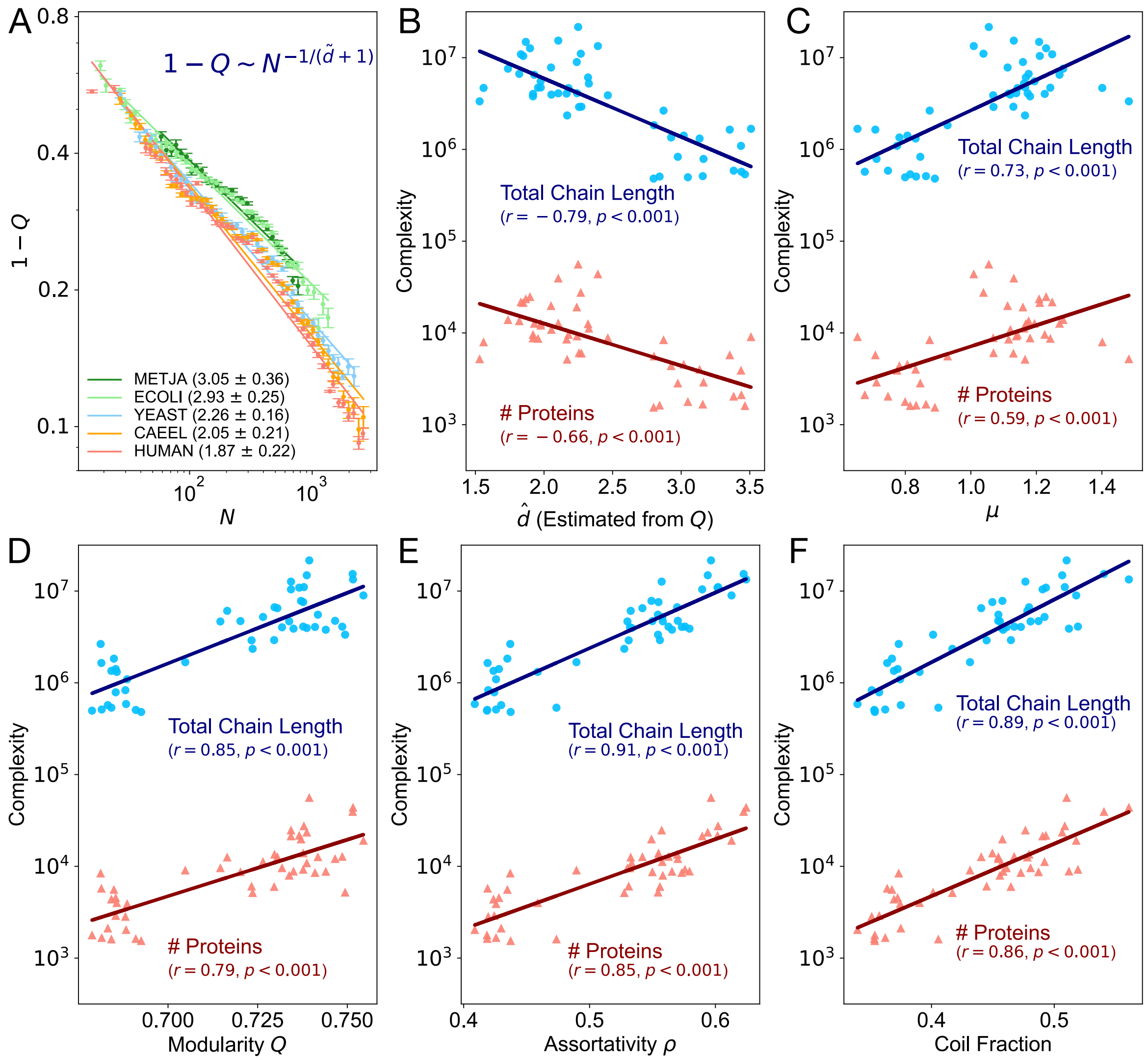


**Figure S3. Structural and topological analyses suggest a decreasing fractal dimension of constituent proteins as the organismal complexity increases.** (A) For the proteins from five selected organisms, the size (chain length $N$) dependence of $1 - Q$, where $Q$ denotes the modularity of the residue contact network. Here, for the data in all the bins, the average standard error of $Q$ is below 1% of the mean value. The estimated fractal dimension $\tilde{d}$ is obtained by robust fittings with Theil–Sen estimators (95% confidence interval). (B-F) For the 48 organisms in AlphaFold DB, the measures of organismal complexity (the total number of proteins and the total chain length of the proteins in the organism proteome) vs. (B) the fitted fractal dimension $\tilde{d}$ and (C) scaling coefficient $\mu$ (estimated from the scaling relation $\lambda_1 \sim N^{-\mu}$). For the proteins with chain lengths $N \approx 250$ from 48 organisms in AlphaFold DB, the measures of organismal complexity vs. the mean values of (D) modularity $Q$, (E) assortativity $\rho$, and (F) coil fractions (calculated by DSSP algorithm). The results suggest that the increasing organismal complexity is accompanied by a decreasing fractal dimension, slower vibration, and higher flexibility of constituent proteins. Moreover, it is observed that higher organismal complexity is associated with higher modularity and higher assortativity of the residue contact network of the constituent proteins.

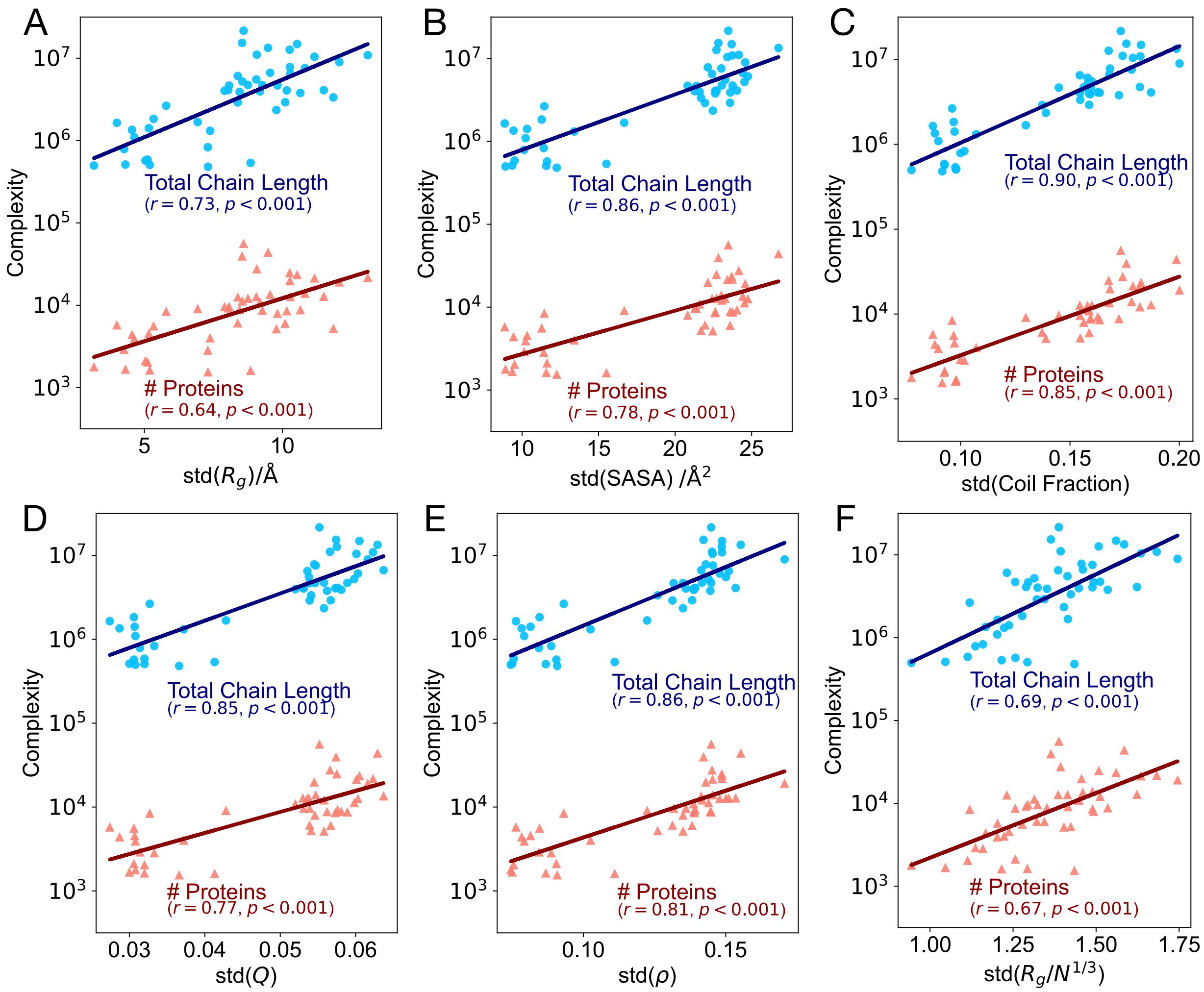


**Figure S4. The correlation between the structural diversity of constituent proteins and the complexity of organisms.** For the proteins with chain lengths $N \approx 250$ from 48 organisms in AlphaFold DB, the measures of organismal complexity (the total number of proteins and the total chain length of the proteins in the organism proteome) vs. the standard deviation of (A) radius of gyration $R_g$, (B) solvent-accessible surface area (SASA) per residue, (C) coil fraction, (D) modularity $Q$, and (E) assortativity $\rho$. (F) For all the proteins from the proteomes of 48 organisms in AlphaFold DB, the measures of organismal complexity vs. standard deviation of the normalized $R_g$ (i.e., $R_g/N^{1/3}$). These results show that the structural diversity of constituent proteins also correlates with organismal complexity.

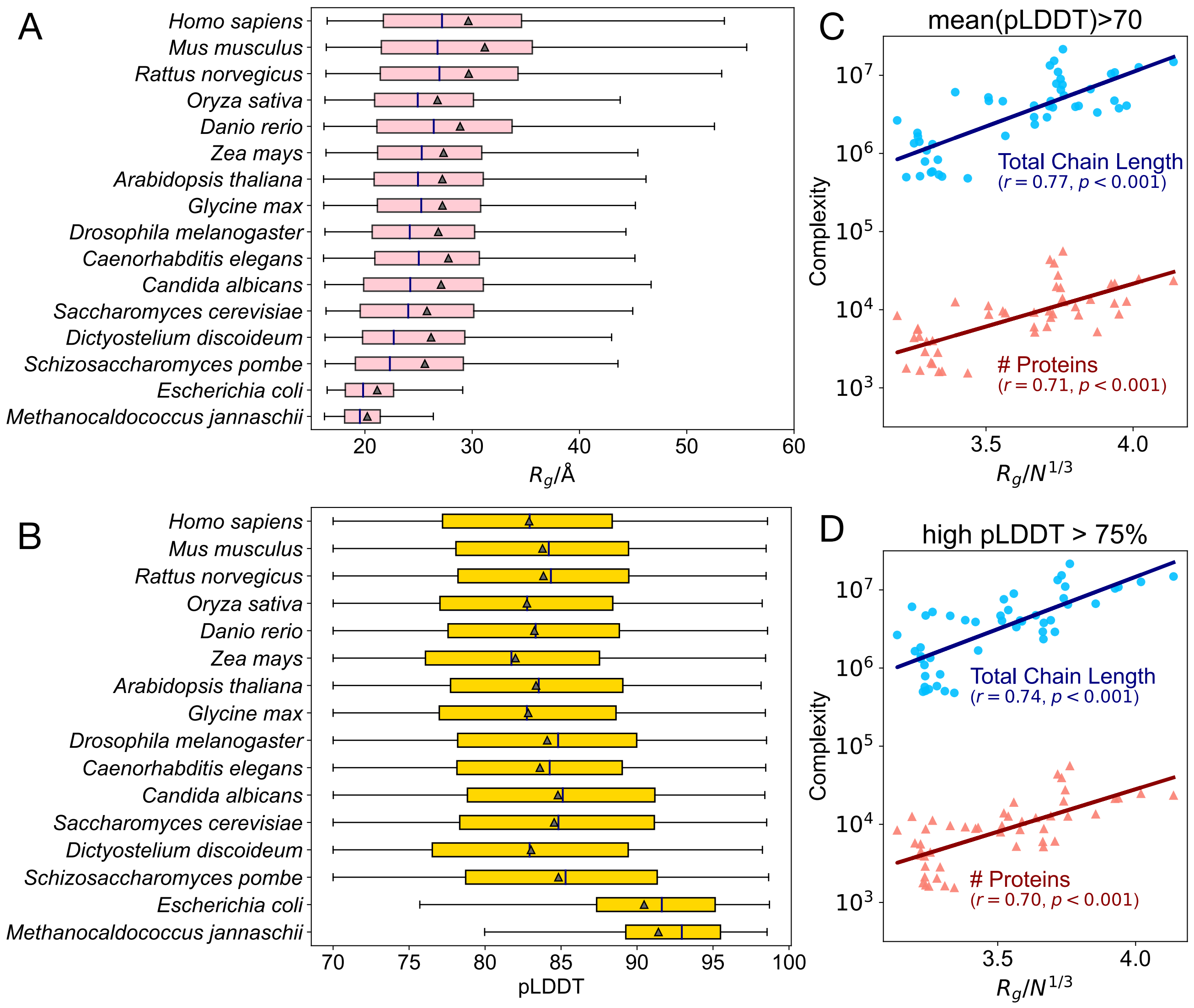


**Figure S5. The correlation between the flexibility of constituent proteins and organismal complexity is robust to the selection of the confidence threshold of structure prediction.** (A) When we filter the AlphaFold-predicted structures with average residue pLDDT ≤ 70, for the proteins with chain lengths $N \approx 250$ ($225 \leq N < 275$), the distributions of $R_g$ are shown as the box-and-whiskers (extreme values not shown). A similar correlation (as shown in Fig. 1A) can also be observed. (B) For the proteins in subplot (A), the distribution of their average pLDDT. It is observed that, compared with prokaryotic proteins, the eukaryotic proteins have a lower average pLDDT. Such a result also indicates higher flexibility of constituent proteins is correlated with higher organismal complexity. In subplots (A) and (B), the orderings of the organisms are based on Fig. 1A. (C-D) For the 48 organisms in AlphaFold DB, when we select the proteins with relatively higher pLDDT from the proteome, the correlation between the measures of organismal complexity and median normalized $R_g$ (i.e., $R_g/N^{1/3}$) still hold. In subplot (C), only the predicted structures with average residue pLDDT > 70 are selected. In subplot (D), only those predicted structures with 75% of their residues having pLDDT >70 are selected. These results further validate that our observations in the Main Text also hold for proteins with relatively stable native structures.

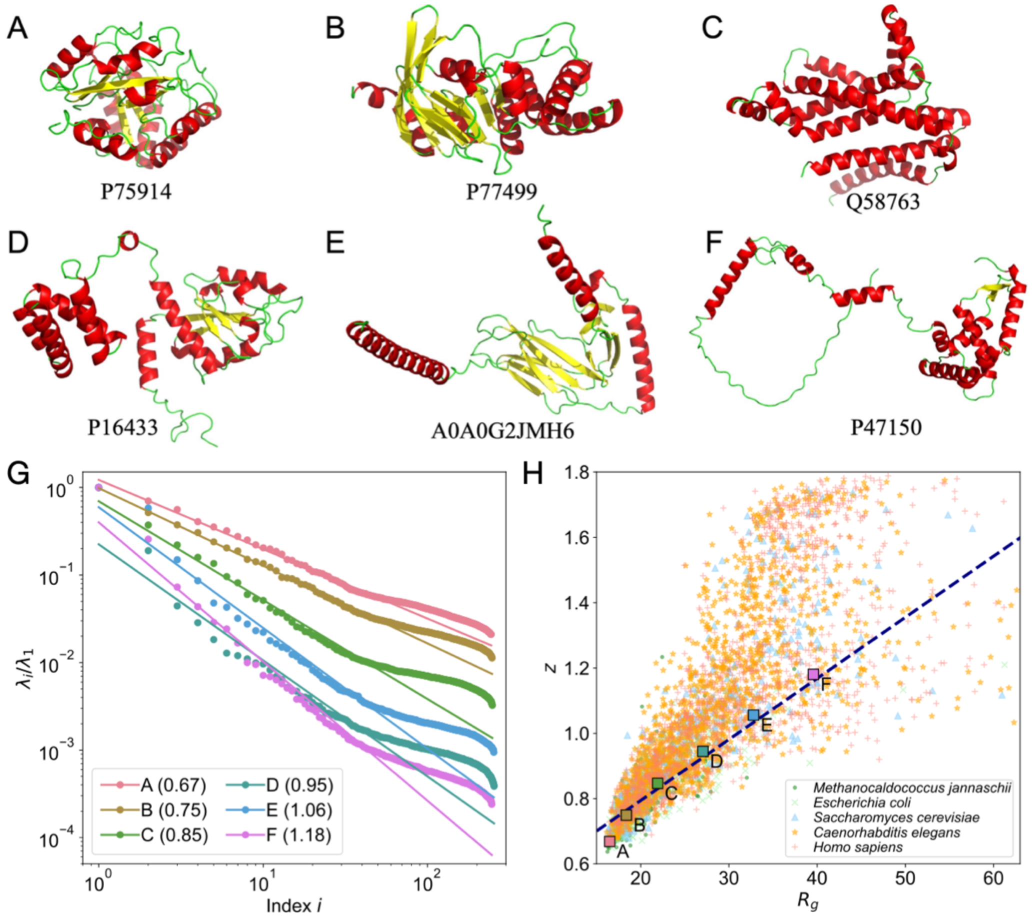


**Figure S6. The proteins with similar chain lengths may have different structures and corresponding vibration spectra.** (A-F) The cartoon illustrations of the six selected proteins in our dataset. Their UniProt codes are also listed in the figure. (G) For the proteins illustrated above, the rank-size distributions of the vibration spectra and the power-law fitting results are shown. The fitted Zipf's coefficient $z$'s are listed in the legend. Here, the fittings are based on the top 25% of the eigenvalues. (H) For the proteins with similar chain lengths $N \approx 250$ $(225 \leq N < 275)$ from the five selected organisms, the scattering plot of their $z$ vs. $R_g$. The trend line in navy shows that Zipf's coefficient $z$ is positively correlated with $R_g$. The crosses in the subplot mark the selected proteins illustrated above.

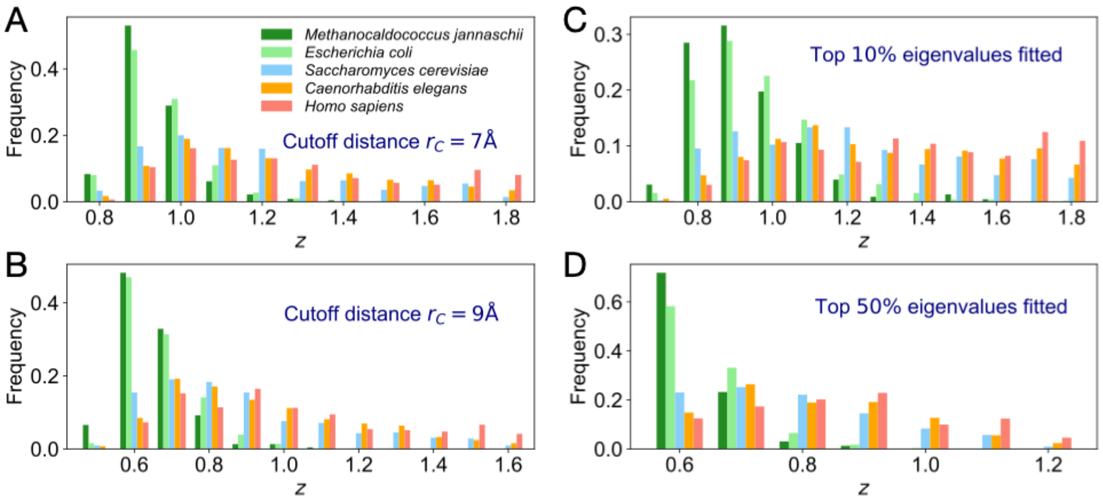


**Figure S7. The correlation between organismal complexity and the Zipf's coefficients of the constituent proteins is robust to the different calculations or fitting methods.** (A-D) With different calculations or fitting details (as indicated within the figure), the obtained distributions of Zipf's exponent *z* for proteins with similar chain lengths in the five selected organisms. In the normal mode analysis based on the Gaussian network model, the selection of the cutoff distance (A) $r_C = 7$ Å and (B) $r_C = 9$ Å may lead to the variations in the values of the fitted *z*, but the correlation between organismal complexity and the Zipf's coefficients is robust to the selection of $r_C$. With $r_C = 8$ Å, in fitting the rank-size distributions of the vibration spectra, the selection of top (C) 10% and (D) 50% also may lead to variations in fitted *z*, but correlation between organismal complexity and the Zipf's coefficients is robust to the selected number of eigenvalues.

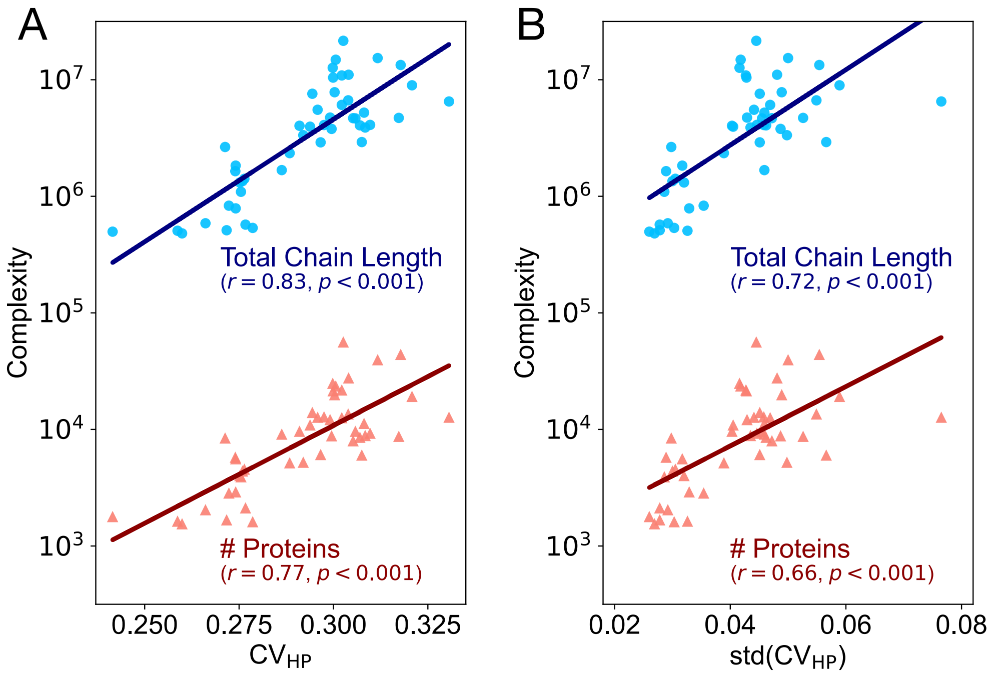


**Figure S8. Organismal complexity correlates with the hydrophilic-hydrophobic segregation in protein sequences.** For the 48 organisms in AlphaFold DB, the measures of organismal complexity (the total number of proteins and the total chain length of the proteins in the organism proteome) vs. (A) the average hydropathy variation $CV_{HP}$, and (B) the standard deviation of $CV_{HP}$. The correlations shown in subplot (A) indicate that organismal complexity is correlated with hydrophilic-hydrophobic segregation in the sequence of the constituent proteins. The correlations shown in subplot (B) suggest that organismal complexity is also correlated with the diversity in the sequences of constituent proteins.

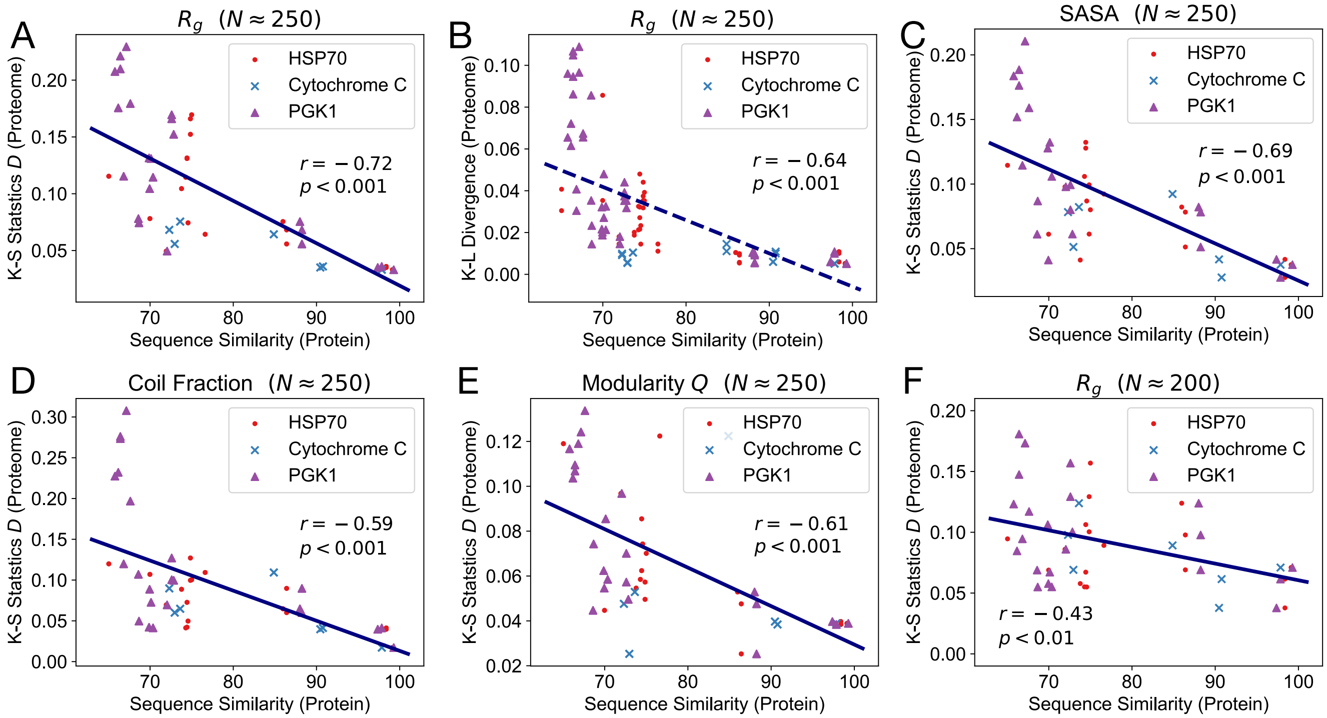


**Figure S9. Sequence similarity of homologous proteins correlates with the statistical structural similarity of the proteomes.** (A-E) For proteins with chain lengths $N \approx 250$ from the proteomes of different organisms, the correlations between statistical structural similarity (with different measures) and sequence similarity are illustrated. In all the subplots, the sequences of the heat shock protein (HSPA9, HomoloGene: 39452), cytochrome C (CYC1, HomoloGene:55617), and phosphoglycerate kinase 1 (PGK1, HomoloGene:21542) are taken as examples for analysis. The phylogenetical distance between two sequences is measured as the sequence similarity based on Clustal Omega. In the comparisons, only the sequences with similarities higher than 65% are considered. Meanwhile, based on the structural statistics of the constituent proteins, the statistical similarity between two proteomes can be calculated. (A) The statistical structural similarity is measured as the KS statistics $D$ between the two $R_g$ distributions and (B) as the Kullback–Leibler (KL) divergence between the two $R_g$ distributions. Since the KL divergence between two distributions is non-symmetric, there are more data points in this subplot (B). The statistical structural similarity is measured (C) as the KS statistics $D$ between the two distributions of SASA, (D) as the KS statistics $D$ between the two distributions of coil fractions, and (E) as the KS statistics $D$ between the two distributions of modularity $Q$. (F) For proteins with chain lengths $N \approx 200$ from the proteomes of different organisms, the statistical structural similarity is measured as the KS statistics $D$ between the two $R_g$ distributions. The results in the subplots indicate that the lineages with high sequence similarity also show high statistical structural similarity.

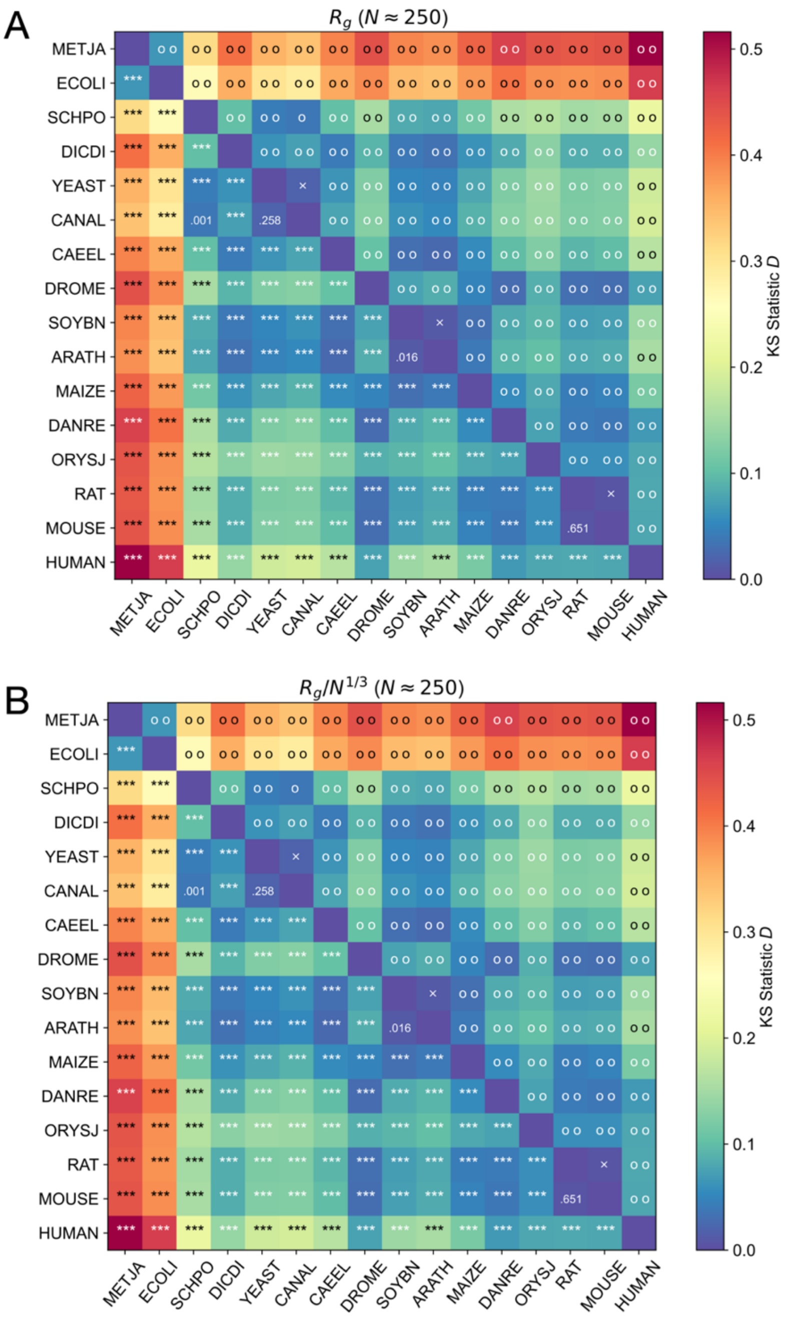


**Figure S10. The two-sample KS test results for the 16 model organisms.** (A) For proteins with similar chain lengths $N \approx 250 (225 \leq N < 275)$ from the 16 model organisms, the KS test results of the $R_g$ distributions. (B) For the proteins (with different chain lengths) from the 16 model organisms, the KS test results of the normalized $R_g$ (i.e., $R_g/N^{1/3}$) distributions. In subplots (A) and (B), the color of each entry shows the KS statistics $D$, and the numbers in the lower triangle show the corresponding $p$-values (where *** denotes $p < 0.001$). The entries in the upper triangle show whether the null hypothesis (both samples come from a population with the same distribution) can be rejected. Here, one circle denotes that the null hypothesis can be rejected at level $\alpha = 0.01$, (i.e.,$\tau(\alpha = 0.01) \geq 1$, but $\tau(\alpha = 0.001) < 1$), two circles denotes that the null hypothesis can be rejected at level $\alpha = 0.001$ (i.e., $\tau(\alpha = 0.001) \geq 1$), and the cross denotes that the null hypothesis cannot be rejected at level $\alpha = 0.01$ (i.e., $\tau(\alpha = 0.01) < 1$).

**Table S1. Detailed information of the dataset used in this study.** The species names, codes, reference proteome, the number of the protein structures, and the total number of residues are listed. The database includes 564,449 predicted protein structures from the proteomes of 48 organisms.

### (A) Model Organisms

| | Species | Common Name | Species Code | Reference Proteome | # Proteins | # Residues |
|---|---|---|---|---|---|---|
| 1 | *Arabidopsis thaliana* | *Arabidopsis* | ARATH | UP000006548 | 27,434 | 11,038,195 |
| 2 | *Caenorhabditis elegans* | Nematode worm | CAEEL | UP000001940 | 19,694 | 7,812,271 |
| 3 | *Candida albicans* | *C. albicans* | CANAL | UP000000559 | 5,974 | 2,918,981 |
| 4 | *Danio rerio* | Zebrafish | DANRE | UP000000437 | 24,664 | 12,660,481 |
| 5 | *Dictyostelium discoideum* | *Dictyostelium* | DICDI | UP000002195 | 12,622 | 6,507,876 |
| 6 | *Drosophila melanogaster* | Fruit fly | DROME | UP000000803 | 13,458 | 6,657,130 |
| 7 | *Escherichia coli* | *E. coli* | ECOLI | UP000000625 | 4,363 | 1,349,576 |
| 8 | *Glycine max* | Soybean | SOYBN | UP000008827 | 55,799 | 21,578,290 |
| 9 | *Homo sapiens* | Human | HUMAN | UP000005640 | 23,391 | 14,850,259 |
| 10 | *Methanocaldococcus jannaschii* | *M. jannaschii* | METJA | UP000000805 | 1,773 | 497,291 |
| 11 | *Mus musculus* | Mouse | MOUSE | UP000000589 | 21,615 | 10,884,613 |
| 12 | *Oryza sativa* | Asian rice | ORYSJ | UP000059680 | 43,649 | 13,377,036 |
| 13 | *Rattus norvegicus* | Rat | RAT | UP000002494 | 21,272 | 10,429,766 |
| 14 | *Saccharomyces cerevisiae* | Budding yeast | YEAST | UP000002311 | 6,040 | 2,902,659 |
| 15 | *Schizosaccharomyces pombe* | Fission yeast | SCHPO | UP000002485 | 5,128 | 2,345,836 |
| 16 | *Zea mays* | Maize | MAIZE | UP000007305 | 39,299 | 15,342,138 |

### (B) Organisms related to global health

| | Species | Common Name | Species Code | Reference Proteome | # Proteins | # Residues |
|---|---|---|---|---|---|---|
| 17 | *Ajellomyces capsulatus* | Ajellomyces capsulatus | AJECG | UP000001631 | 9,199 | 4,097,952 |
| 18 | *Brugia malayi* | Brugia malayi | BRUMA | UP000006672 | 8,743 | 3,786,006 |
| 19 | *Campylobacter jejuni* | *C. jejuni* | CAMJE | UP000000799 | 1,620 | 506,322 |
| 20 | *Cladophialophora carrionii* | Cladophialophora carrionii | 9EURO1 | UP000094526 | 11,170 | 5,219,978 |
| 21 | *Dracunculus medinensis* | Dracunculus medinensis | DRAME | UP000274756 | 10,834 | 3,956,557 |
| 22 | *Enterococcus faecium* | Enterococcus faecium | ENTFC | UP000325664 | 2,823 | 830,755 |
| 23 | *Fonsecaea pedrosoi* | Fonsecaea pedrosoi | 9EURO2 | UP000053029 | 12,509 | 6,083,143 |
| 24 | *Haemophilus influenzae* | *H. influenzae* | HAEIN | UP000000579 | 1,662 | 511,443 |

| | | | | | | |
|---|---|---|---|---|---|---|
| 25 | *Helicobacter pylori* | *H. pylori* | HELPY | UP000000429 | 1,538 | 480,731 |
| 26 | *Klebsiella pneumoniae* | *K. pneumoniae* | KLEPH | UP000007841 | 5,727 | 1,643,098 |
| 27 | *Leishmania infantum* | *L. infantum* | LEIIN | UP000008153 | 7,924 | 4,678,633 |
| 28 | *Madurella mycetomatis* | Madurella mycetomatis | 9PEZI1 | UP000078237 | 9,561 | 4,660,506 |
| 29 | *Mycobacterium leprae* | Mycobacterium leprae | MYCLE | UP000000806 | 1,602 | 535,547 |
| 30 | *Mycobacterium tuberculosis* | *M. tuberculosis* | MYCTU | UP000001584 | 3,988 | 1,313,991 |
| 31 | *Mycobacterium ulcerans* | Mycobacterium ulcerans | MYCUL | UP000020681 | 9,033 | 1,677,574 |
| 32 | *Neisseria gonorrhoeae* | *N. gonorrhoeae* | NEIG1 | UP000000535 | 2,106 | 571,507 |
| 33 | *Nocardia brasiliensis* | Nocardia brasiliensis | 9NOCA1 | UP000006304 | 8,372 | 2,648,670 |
| 34 | *Onchocerca volvulus* | Onchocerca volvulus | ONCVO | UP000024404 | 12,047 | 4,735,343 |
| 35 | *Paracoccidioides lutzii* | Paracoccidioides lutzii | PARBA | UP000002059 | 8,794 | 3,890,384 |
| 36 | *Plasmodium falciparum* | *P. falciparum* | PLAF7 | UP000001450 | 5,187 | 3,343,123 |
| 37 | *Pseudomonas aeruginosa* | *P. aeruginosa* | PSEAE | UP000002438 | 5,556 | 1,833,487 |
| 38 | *Salmonella typhimurium* | *S. typhimurium* | SALTY | UP000001014 | 4,526 | 1,413,095 |
| 39 | *Schistosoma mansoni* | Schistosoma mansoni | SCHMA | UP000008854 | 13,865 | 7,581,408 |
| 40 | *Shigella dysenteriae* | *S. dysenteriae* | SHIDS | UP000002716 | 3,893 | 1,093,053 |
| 41 | *Sporothrix schenckii* | Sporothrix schenckii | SPOS1 | UP000018087 | 8,652 | 4,705,197 |
| 42 | *Staphylococcus aureus* | *S. aureus* | STAA8 | UP000008816 | 2,888 | 787,862 |
| 43 | *Streptococcus pneumoniae* | *S. pneumoniae* | STRR6 | UP000000586 | 2,030 | 587,055 |
| 44 | *Strongyloides stercoralis* | Strongyloides stercoralis | STRER | UP000035681 | 12,613 | 5,524,754 |
| 45 | *Trichuris trichiura* | Trichuris trichiura | TRITR | UP000030665 | 9,564 | 4,027,259 |
| 46 | *Trypanosoma brucei* | Trypanosoma brucei | TRYB2 | UP000008524 | 8,491 | 4,062,090 |
| 47 | *Trypanosoma cruzi* | *T. cruzi* | TRYCC | UP000002296 | 19,036 | 8,959,643 |
| 48 | *Wuchereria bancrofti* | Wuchereria bancrofti | WUCBA | UP000270924 | 12,721 | 4,087,813 |

**Table S2. The Kolmogorov–Smirnov (KS) test for the distributions of the proteins from the five selected organisms.** For the proteins with similar chain lengths $N \approx 250$ $(225 \leq N < 275)$ in different species, we compare their distributions of the (A) radii of gyration $R_g$, (B) Coil fraction, and (C) hydropathy variation $CV_{HP}$. The KS statistic $D$ and the corresponding $p$-values are listed in the upper and lower triangles of the tables, respectively. Here, *** denotes the $p$-values below 0.001. These results show that, when we compare the structures (radii of gyration or secondary structure) of proteins with similar chain lengths, there is no significant difference ($p > 0.01$) between *M. jannaschii* (archaea) and *E. coli* (bacteria). In contrast, when comparing their sequences ($CV_{HP}$), there is a significant difference between *M. jannaschii* and *E. coli*, while the hydropathy segregation in the protein sequences of the eukaryote organisms do not differ significantly ($p > 0.01$) from each other.

**(A)**

| $R_g$ | METJA | ECOLI | YEAST | CAEEL | HUMAN |
|---|---|---|---|---|---|
| METJA | | 0.0946 | 0.4657 | 0.5598 | 0.6288 |
| ECOLI | 0.1104 | | **0.3975** | 0.4776 | 0.564 |
| YEAST | *** | *** | | 0.1154 | 0.2076 |
| CAEEL | *** | *** | *** | | 0.166 |
| HUMAN | *** | *** | *** | *** | |

**(B)**

| Coil % | METJA | ECOLI | YEAST | CAEEL | HUMAN |
|---|---|---|---|---|---|
| METJA | | 0.1158 | 0.3622 | 0.4128 | 0.5335 |
| ECOLI | 0.0265 | | 0.2754 | 0.3429 | 0.4607 |
| YEAST | *** | *** | | 0.1199 | 0.2274 |
| CAEEL | *** | *** | *** | | 0.1271 |
| HUMAN | *** | *** | *** | *** | |

**(C)**

| $CV_{HP}$ | METJA | ECOLI | YEAST | CAEEL | HUMAN |
|---|---|---|---|---|---|
| METJA | | 0.4962 | 0.5956 | 0.6217 | 0.6437 |
| ECOLI | *** | | 0.2056 | 0.2598 | 0.2694 |
| YEAST | *** | *** | | 0.0683 | 0.0745 |
| CAEEL | *** | *** | 0.0895 | | 0.0364 |
| HUMAN | *** | *** | 0.0525 | 0.291 | |
